# Structure-function analysis of PI3K signalling cascade base editing screens in cancer cells

**DOI:** 10.64898/2026.09.08.748517

**Authors:** Katrina McCarten, Barbara Abreu, Benoit Baillif, Lisa Koob, Jong Sook Ahn, Cansu Dincer, Julian Esselborn, Emre Karakoc, R. Frederick Ludlow, John Lyons, Gabriele Picco, Mamta Sharma, Marcel L. Verdonk, Mark Wade, Samantha Walker, Alexander Watterson, Matthew A. Coelho, Thomas G. Davies, Mathew J. Garnett

## Abstract

Knowledge of protein structure and function underpins rational drug discovery, yet many targets lack known selectively druggable sites. Furthermore, the identification of secondary druggable sites offers a strategy to overcome drug resistance. Fragment-based drug discovery (FBDD) can identify new ligandable binding pockets, though how to triage those with the ability to exert biologically relevant effects can be unclear. Systematic approaches to identify novel functionally-important protein sites for therapeutic intervention, such as allosteric pockets or protein–protein interaction (PPI) interfaces, has the potential to accelerate drug discovery, particularly when combined with structure-based hit-finding modalities. The phosphoinositide 3-kinase (PI3K) signalling pathway is frequently altered in human cancer and resistance to approved inhibitors is an ongoing challenge. Here, we performed large-scale CRISPR base editing mutagenesis screens across 30 PI3K pathway proteins in three disease-relevant cancer cell models to systematically map functional residues. Integration of base editing data with structural information identified residues corresponding to known catalytic sites, fragment-binding pockets and PPI interfaces, providing validation for the approach. Additionally, we identified putative allosteric pockets near regions of undefined function. Together, these findings establish high-throughput base editing mutagenesis combined with structural analysis as a scalable strategy to delineate structure–function relationships and inform drug development.

## Introduction

Detailed knowledge of protein function and structure can facilitate drug discovery. Although sites of enzymatic activity are the most common point of intervention by small molecule therapeutics, many targets lack a precedented and druggable site that can be selectively targeted. In these cases, it may be desirable to modulate activity by targeting an allosteric pocket or a region involved in a protein–protein interaction (PPI). Targeting a specific PPI could also enable development of therapeutics targeting specific protein functions and effector pathways, while sparing others, potentially reducing safety concerns where complete ablation of function is undesirable^1^. Moreover, alternative approaches to re-target established drug targets can overcome resistance, as evidenced by the clinical success of second and third generation EGFR and ABL inhibitors.

State-of-the-art approaches for interrogating protein function and structure in drug design include high-resolution structural techniques, as well as functional mapping strategies like saturation genome editing (SGE) and deep mutational scanning (DMS)^2,3^. Structural methods require purified proteins and are limited to the acquisition of static snapshots outside of cellular contexts, whereas SGE and DMS can be performed in cellular contexts but remain challenging to scale across multiple proteins and physiologically relevant cellular models. Unlike CRISPR tiling which creates random indels^4^, CRISPR base editing can introduce precise base-pair C→T and A→G alterations in diverse cell types in a high-throughput manner, allowing the introduction of many mutations to the endogenous protein in a single assay^5,6^. This approach allows a route to probing protein structure–function relationships and potential points of intervention, at scale, in physiologically relevant cell types.

The phosphoinositide 3-kinase (PI3K) pathway is central to integrating signals controlling cellular growth, proliferation and survival, and is among the most frequently deregulated pathways in cancer^7^. Approximately one third of human tumours harbour genomic alterations in the PI3K pathway^8^ and these alterations are particularly enriched in breast, gastrointestinal, head and neck and gynaecological cancers^9^. Activating variants in *PIK3CA* occur in ∼14% of cancers, while loss-of-function alterations in *PTEN* (∼8%) and *PIK3R1* (∼4%) additionally drive pathway hyperactivation^9^. The prevalence of these alterations has driven extensive efforts to therapeutically target PI3K, mTOR and AKT. Despite the generation of multiple classes of compounds, including PI3K isoform-selective agents with improved toxicity profiles, most clinically approved inhibitors target conserved catalytic domains^10–12^. The emergence of resistance in patients towards PI3K inhibitors, particularly through PTEN loss and AKT1 activation, remains a clinical challenge^10,13^. A systematic approach to define functionally critical and targetable residues across the pathway is therefore highly desirable.

In this proof-of-concept study, functional sites on PI3K pathway proteins in disease-relevant cancer cell lines were mapped using CRISPR base editing mutagenesis combined with structural interpretation. We validated this approach by identifying known druggable sites on target proteins and identified putative targetable allosteric pockets. This study highlights the ability of our platform to uncover novel sites offering a new complementary strategy to enable drug discovery.

## Results

### Systematic mutagenesis of the PI3K signalling pathway

To construct structure–function maps, we performed fitness-based CRISPR base editing dropout screens across 30 genes of the PI3K pathway, prioritising genes with available structural information (Figure 1a and 1b). *PIK3CA* mutant cancer cell lines MCF7 (breast), T47D (breast) and HGC27 (gastric) were engineered to stably express doxycycline-inducible CBE (BE3.9max-Cas9NGN^14^) or ABE (ABE8e-Cas9NGN^15^). Clonally-derived lines with the highest base editing efficiency were selected using the BE-FLARE or GFP stop codon reporter constructs for CBE and ABE, respectively^16,17^ (Supplementary Figure 1a). All three models display dependency on PI3K pathway genes, supporting their use for viability-based screening^18^.

**Figure 1:**
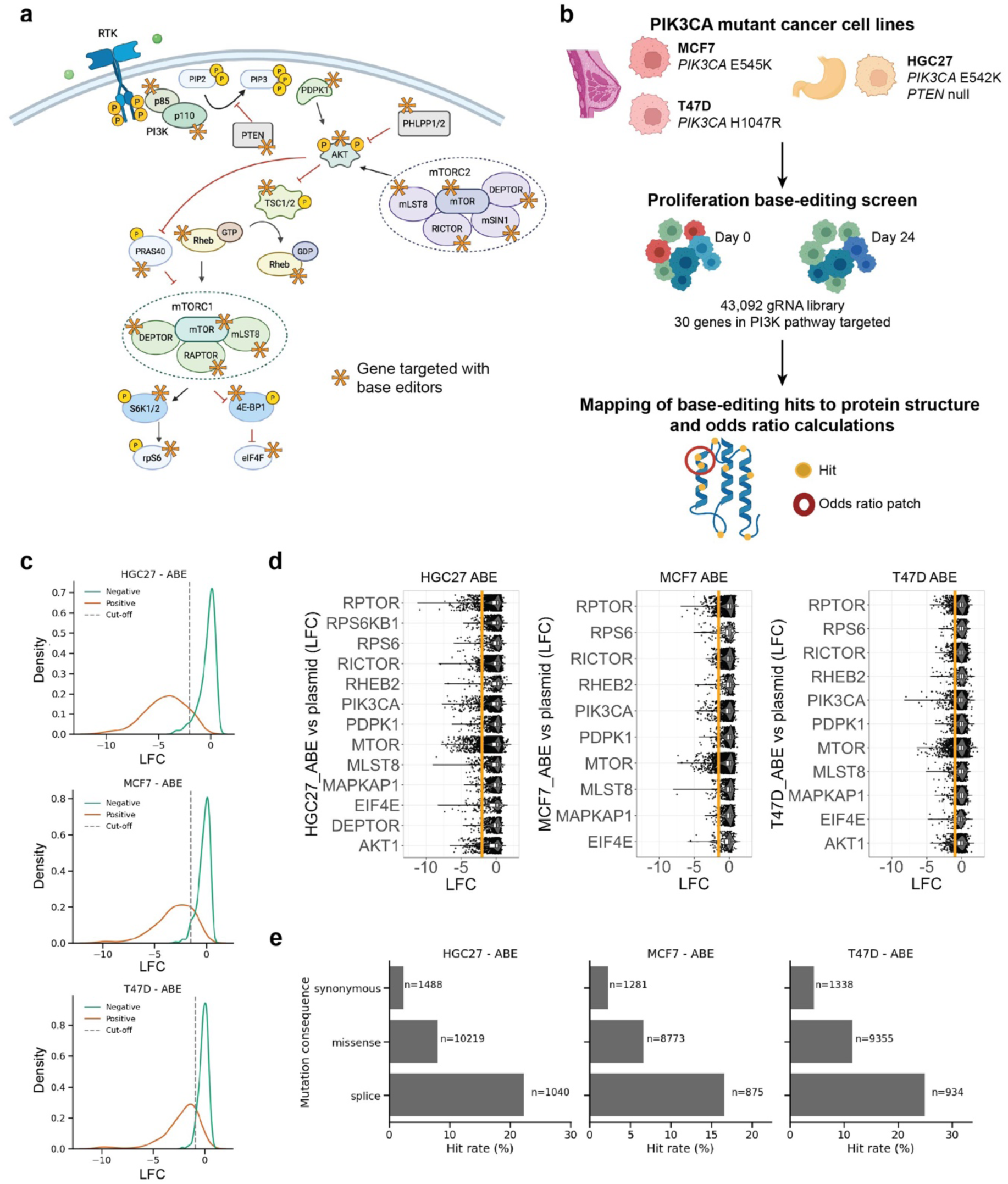
Mutagenesis screen of the PI3K signalling pathway. **a**) PI3K pathway depiction. Activation shown by black arrows and inhibition by red lines. The proteins encoded by the genes targeted in the CRISPR base editing screen are highlighted with asterisks. Schematic created with BioRender.com. **b**) Schematic of base editing screens in PI3K mutant cancer cell lines to find variants with a negative impact on proliferation/survival-based screens. Mapping onto the 3D structure of a protein with an odds ratio approach allows the selection of the most impactful regions. **c**) Distribution plots of internal positive and negative screen control guides for ABE screens in HGC27, MCF7 and T47D cell lines. The dashed line indicates the LFC value cut off with a 5% FDR for hit calling. **d**) Violin plot showing the depletion of guides across all essential genes within the PI3K pathway for ABE in HGC27, MCF7 and T47D cells. Boxplots show the median and interquartile range. The LFC cut off for depleted guides is marked by an orange line. The essentiality for each gene was determined from DepMap data^21^, in which all genes with gene effect lower than -0.5 were considered essential. **e**) Distribution of percentage of hits by most severe predicted edit type per guide for essential genes listed in (d). The number of guides reported differs between cell lines because some genes are essential only in certain cell lines. ABE: Adenine Base Editor; FDR: False Discovery Rate; LFC: Log_2_ Fold Change

Using BEstimate^19^, we designed a pooled library of 43,092 single guide RNAs (sgRNAs) to target genes based on all available Protospacer Adjacent Motif (PAM) sites, supplemented with control sgRNA for essential and non-essential genes (Supplementary Table 1 and 2). Screens were performed in duplicate. Replicate screens correlated well, with better correlations observed for ABE than for CBE, the latter reflecting reduced effect sizes likely arising from a lower base editing efficiency (Supplementary Figure 1b and 1c).

As expected, positive control sgRNA (splice sites of essential genes^20^) showed negative fitness scores (log fold changes < 0), whereas negative control guides (intergenic, non-targeting and splice sites of non-essential genes^20^) had little or no effect and were centred around zero (Figure 1c and Supplementary Figure 2a). However, separation between positive and negative controls was diminished in CBE versus ABE, and owing to poor discriminatory power, we excluded the T47D–CBE screens from downstream analyses.

To robustly identify sgRNAs associated with loss of cell fitness, we established a data-driven threshold for each cell line base editor combination based on the log-fold change (LFC) effect size and applying a 5% false discovery rate (FDR) (see methods). Guides with a LFC below this threshold were labelled as depleted (Figure 1d), resulting in strong enrichment of positive control guides among depleted sgRNAs. For example, for HGC27–ABE, the recovery rate of negative controls called as depleted is 4.5%, compared with 90% among positive controls. Across genes within each cell line, the proportion of depleted guides correlated well with effect size of the gene knockout (range: -0.48 for HGC27–CBE to -0.84 for MCF7–ABE; Supplementary Figure 3b and 3d). For instance, in HGC27–ABE screens, the proportion of depleted guides ranged from 0.9% for the non-essential gene *PTEN*, to 16% for the essential gene *AKT1*.

In total, we identified 4463 sgRNAs called as depleted across all genes and cell lines, with 3053, 2511 and 2332 sgRNAs for HGC27, MCF7, and T47D respectively, with 883 depleted in both CBE cell lines and 970 depleted in all three ABE cell lines. The number of depleted sgRNAs varied considerably on a per gene basis (median=87 sgRNAs; range 26–703 sgRNA), due to variation in gene essentiality in an individual cell line, cell line–specific editor efficiency, and the number of targeting sgRNAs per gene (Figure 1d, Supplementary Figure 2b, 3a and 3c).

Together, these analyses demonstrate the sensitivity and specificity of our base editing screens and validate our ability to discriminate essential from non-essential genes, supporting the robust identification of sgRNAs affecting cell fitness.

### Concordance Between Predicted Variant Effects and Screen Derived Fitness Hits

To further assess screen performance, we compared hit rates among sgRNAs targeting essential genes in the PI3K pathway after stratifying them by predicted mutation consequence (synonymous, missense or splice) annotated using Ensembl Variant Effect Predictor (VEP)^22^ (Figure 1e and Supplementary Figure 2c). Across all ABE screens, sgRNA predicted to induce splice site disrupting mutations showed the highest hit rates (16.6−25%), followed by those generating missense mutations (6.6−11.5%). A significant fraction of missense hits are predicted to induce leucine-to-proline mutations in ABE screens; this change was associated with the highest hit rate in the screens (Supplementary Figure 4). This is expected, as proline mutations introduce α-helices kinks and backbone rigidification, which ultimately compromises both structure and function^23^. In contrast, guides predicted to introduce synonymous edits exhibited the lowest hit rates (2.3−4.3%). The impact of a sgRNA edit is predicted based on a set base editing window (4−9nt). The exact editing window is known to vary at the individual guide level and some hits within the predicted synonymous category may reflect this^24^. A similar but weaker trend was observed in CBE for HGC27 (3.6 %, 5.9% and 14.0%), and MCF7 (2.9%, 3.4% and 6.5%), for synonymous, missense and splice/stop, respectively. These trends are consistent with the expected functional impact of different mutation classes.

Together, these results demonstrate that our base editing screens capture the expected spectrum of functional effects, providing confidence in their ability to resolve biologically meaningful variation.

### Connecting Variant Function to Protein Structure

Having validated our ability to systematically screen for variant effects, we next examined hit sgRNAs targeting essential PI3K pathway genes in the context of their coding sequence (CDS), Uniprot family, domain annotations footprint, and protein 3D structure.

Figure 2 exemplifies this analysis on p110α and mTOR proteins, while the analyses of other essential genes are displayed in Supplementary Figure 5 and 6. By normalizing the log fold change values per dataset, it is possible to obtain z-scores, which allow the direct comparison of the adenine and cytidine base editors with regard to their effects on viability. Some cell line–specific effects were observed as expected, due to technical variation and cell line–specific gene essentiality. Generally, while most sgRNAs had no effect, clusters of deleterious guides were observed across protein domains, including within key catalytic and functional domains.

**Figure 2:**
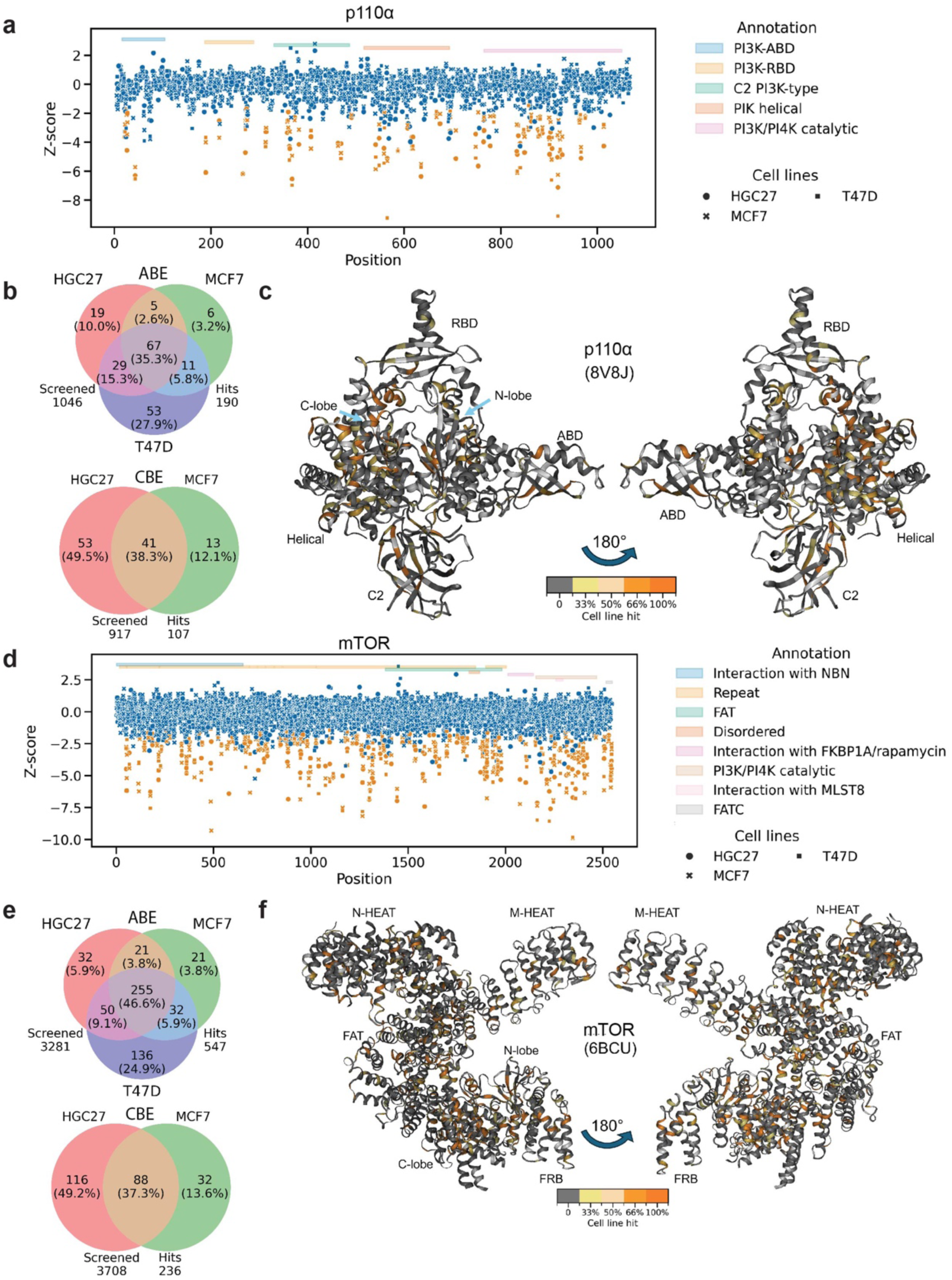
Projection of base editing screens onto p110α and mTOR protein 3D structures. **a**) Z-score for each missense-inducing sgRNA reported on the amino acid residue position for p110α. Points coloured in orange highlight guides that are a hit in all three tested cell lines in ABE, or a hit in both HGC27 and MCF7 in CBE screens. **b**) Venn diagram of the hits targeting *PIK3CA* with percentage of hits shared between HGC27, MCF7 and T47D cell lines in ABE screens (top) and between HGC27 and MCF7 cell lines in CBE screens (bottom) shown. **c**) Percentage of cell line hits for each residue shown on the p110α protein structure (PDB code: 8V8J). **d**) Z-score for each missense-inducing guide reported on the amino acid residue position for mTOR. **e**) Venn diagram of the hits targeting *MTOR* with percentage of hits shared between HGC27, MCF7 and T47D cell lines in ABE screens (top) and between HGC27 and MCF7 cell lines in CBE screens (bottom). f) Percentage of cell line hits for each residue shown on the mTOR protein structure (PDB code: 6BCU). ABE: Adenine base editor; CBE: Cytosine base editor.

Using only guides predicted to produce missense mutations, we designed a systematic approach relying on statistical hypothesis testing to search for protein regions sensitive to mutation across the large number of genes tested. Given a structure or model of a protein, this method generates 3D circular patches of amino acids (Figure 3a), centred around a specific amino acid and including neighbouring residues within a 7 Å radius (approximating the radius of a typical binding-site concavity or small structural region, representing on average 15-20 residues). For each patch, we computed the odds ratio (OR) of robust hits observed in all cell lines (n=3 for ABE and n=2 for CBE) within the patch compared with the remainder of the protein. Henceforth, these patches will be referred as OR patches.

**Figure 3:**
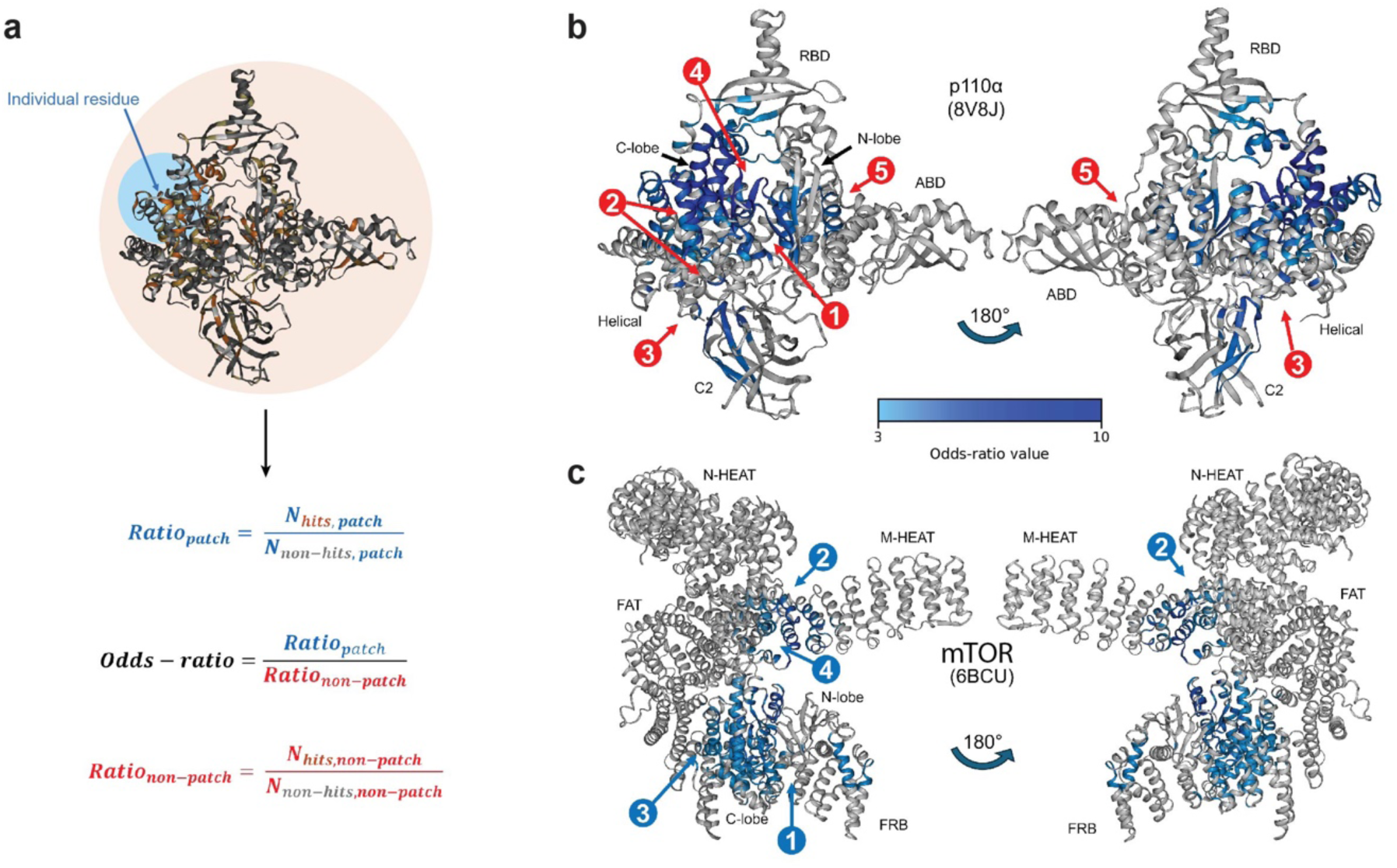
Odds ratio framework for identification of patches within p110α and mTOR protein structures. **a)** For each residue in the protein to be considered a patch centre, an OR patch was defined by examining residues within 7 Å of this central residue. The ratio of hit sgRNAs vs non-hits (coloured in orange, as opposed to non-hits in grey) in a patch (blue in the equation) was calculated as opposed to the ratio in the rest of the protein (red in the equation). This was used to compute the odds-ratio as depicted. Significant OR patches on p110α (**b**) and mTOR (**c**). The ORs are coloured with a light to dark blue scale for ascending values from 3 to 10. Sites selected for further investigation are in red for p110α, and blue for mTOR. ABD: Adaptor-binding domain; FAT: FKBP12 rapamycin-associated protein, ataxia telangiectasia, and transformation or transactivation domain associated protein; FRB: FKBP-rapamycin-binding; OR: Odds ratio; RBD: Ras binding domain.

Significant OR patches were identified across several proteins, including p110α (Figure 3b), mTOR (Figure 3c), Raptor (Supplementary Figure 7a), RICTOR (Supplementary Figure 7b), RPS6 (Supplementary Figure 7c), PDPK1 (Supplementary Figure 7d) and MAPKAP1 (mSIN1) (Supplementary Figure 7e). Here, we highlight a subset of sites comprising one or multiple enriched OR patches in p110α and mTOR for further investigation to validate the approach and to illustrate the types of insights that can be revealed using this method.

### Functional patches within catalytic and allosteric sites

In mTOR and p110α (the catalytic domain of PI3Kα), statistically significant OR patches were found within the kinase region and, most importantly, in the C-lobe, which is responsible for structural stability, substrate binding, and contains the catalytic loop (Supplementary Figure 8a and 8c, sites 1 and 2). In the kinase site of mTOR (Supplementary Figure 8a), we observed multiple guide hits. The hits included guides predicted to mutate multiple conserved residues involved in ATP binding: [V2162A, I2163T], [V2183A, F2184S/L/P], [I2237T], [S2342P] and [Y2225H]. Mutation of many of these amino acids has been associated with resistance to competitive mTOR inhibitors^25^. We also observed hits buried in the C-lobe region of mTOR, which are predicted to give rise to [L2246P], [S2323P, L2324P], [V2326A, M2327T, S2328P], [L2344P, M2345T] and [I2353T]. Predicted mutations in buried structural regions, especially prolines and non-conserved amino acid substitutions with different physiochemical properties, might result in protein unfolding. Hit guides in the catalytic loop predicted to edit [C2361R], [F2371L/P/S], [E2373K] and [F2377L/P/S] were also seen.

In p110α, (Supplementary Figure 8c, site 1, top panel) multiple hits that induce mutations in residues directly involved in ATP binding were identified, including those in conserved residue [Y836C] and mutations [I921V, M922V] and [M922T, V923A] (Supplementary Figure 8d–f). Additionally, we observed two missense hits in the catalytic loop of p110α, which were expected to alter enzymatic activity by inducing non-conservative mutations, [H917R, N918S/G/D] (Supplementary Figure 8d–f) and [N918S/G/D, S919G]. We observed hits in hydrophobic buried regions in the N-lobe of p110α: [L839P, S840P], [L847P]; and in hydrophobic buried regions of the C-lobe: [L877P], [W880R, L881P], [S900L], [Y904H, C905R], [C905R, V906A]. There are also hits targeting buried residues on a strand adjacent to the hinge region ([M833I], [M833T, L834P/S]), which may be important for stabilising the relative orientation of the N- and C-lobes, with the C-lobe being required to localise p110α to the lipid layer, which is needed for PI3Kα-dependent phosphorylation of PIP2 to produce PIP3^26^.

Whilst many of the current drugs/ligands for PI3Kα (the heterodimer formed by the catalytic p110α subunit and regulatory p85α subunits) are orthosteric, allosteric ligands have also been reported^27–29^. In p110α, the C-lobe represents half of the kinase active site (within p110α site 1, Supplementary Figure 8c) but also contains two known allosteric binding sites (within p110α site 2, Supplementary Figure 8c), referred to as top and bottom sites based on their position on the site 2 of Supplementary Figure 8c. The top allosteric site is targeted by an H1047R-selective over wild-type inhibitor^27^, as hypothesised previously^30^, and the bottom allosteric site is targeted by the mutant PI3Kα-selective inhibitors, RLY-2608^28^ and STX-478^29^. p110α site 2 (Supplementary Figure 8c) shows the two allosteric sites opposite the orthosteric site of the C-lobe. The hits [Y904H, C905R] and [C905R, V906A] target residues close to the top allosteric binding site. However, we failed to identify hits located near the bottom allosteric binding site, indicating that while the patch analysis may highlight areas where structure is important, it may also miss some residues that participate in ligand binding because either there are no sgRNAs targeting the region or the available guides do not edit the region appropriately.

Overall, the catalytic sites of mTOR and p110α are visible from clear clusters of hits, and this is reflected by the significant OR patches recovered, validating the approach to identify structural or functional sites of interest. Due to the biological significance of these regions, many drugs and ligands (especially in the case of competitive inhibitors) have been previously associated with these sites.

### OR patches represent functionally relevant protein–protein interfaces

In addition to known catalytic and allosteric sites, multiple OR patches were identified at protein**–**protein interfaces, providing insights that might inform novel drug development approaches. In mTOR, OR site 2 is at a known mTOR–RHEB interface^31^ (Figure 4a). One identified hit on the mTOR FAT domain (FKBP12 rapamycin-associated protein, ataxia telangiectasia, and transformation or transactivation domain associated protein) is [C1303R, W1304R]^32^. Co-competition assays allowed us to validate reduction in growth or viability of MCF7 ABE cells containing the [C1303R, W1304R] edit compared with cells containing a non-essential gene control (Figure 4b and 4c). Western blotting showed that the global levels of mTOR were slightly reduced by the introduction of both C1303R and W1304R mutations and downstream signalling was significantly reduced, as determined by observing lower levels of phosphorylated p70S6K (S6K1) (Figure 4d). Whilst not forming a statistically significant OR patch, we also observed guide hits predicted to generate the [I76T] mutation in ABE screens (233RHEB, Figure 4b) and [D77N] in CBE screens. These residues are on the corresponding face of RHEB that participates in the PPI. Both I76 and D77 are within the switch II region of RHEB and are known to interact with mTOR^33^. Mutation of these residues can impair the ability of GTP-bound RHEB to activate mTORC1^34,35^. Targeting the location of this interaction with a ligand might prevent RHEB binding to mTOR, thereby inhibiting mTORC1 activity.

**Figure 4:**
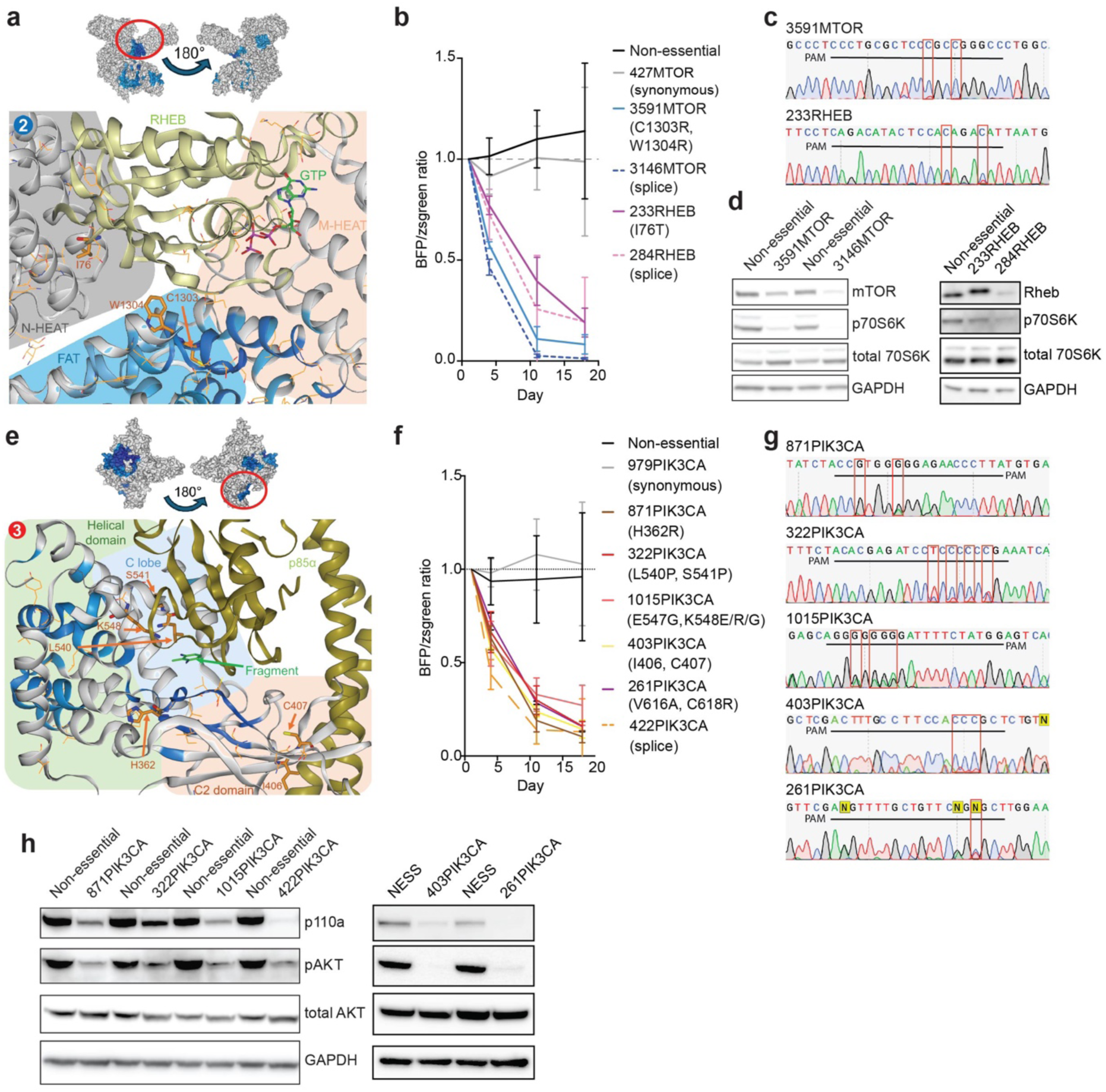
Structural analysis of odds ratio patches on protein–protein interaction sites on p110α and mTOR. **a**) Miniature surface representation of mTOR with indicated OR patch (top). Cartoon representation of the focus on the mTOR–RHEB interface (PDB code: 6BCU; bottom). **b**) Co-competition experiment in MCF7 ABE cells showing the relative proportion of cells expressing tested sgRNAs compared with a non-essential guide control. N=2 for 3591MTOR, 3146MTOR and 284RHEB; N=3 for 427MTOR and 233RHEB. Dashed line at a ratio of 1.0 signals an equal ratio of BFP and zsgreen fluorescence. **c)** Sanger sequencing of MCF7 ABE cells to confirm sgRNA editing of target site. **d**) Western blots showing protein abundance in MCF7 ABE cells transfected with tested sgRNAs at day 7. GAPDH was used as a loading control. **e**) Miniature surface representation of p110α with indicated patch (top). Focus on p110α and p85α interaction, on the phosphopeptide binding site with fragment binding (PDB code: 5SXK; bottom). **f)** Co-competition experiment in MCF7 ABE cells showing relative proportion of MCF7 ABE cells expressing tested sgRNAs compared with a non-essential guide. N=2 for 871PIK3CA, 322PIK3CA, 261PIK3CA and 403PIK3CA; N=3 for 979PIK3CA, 1015PIK3CA and 422PIK3CA. Dashed line at a ratio of 1.0 signals an equal ratio of BFP and zsgreen fluorescence. **g**) Sanger sequencing of MCF7 ABE cells transfected with targeting sgRNAs. **h**) Western blots showing protein abundance in MCF7 ABE cells transfected with tested sgRNAs at day 7.

In p110α, OR site 3 is located between the helical domain and C2 domain of the p110α subunit and the N-terminal Src Homology 2 (nSH2) domain of the p85α subunit (*PIK3R1*) (Figure 4e). This region is also known as the phosphopeptide binding site. Binding of the nSH2 domain of p85α to the C2, helical and kinase domains of p110α stabilises the p110α–p85α interaction such that the inter-SH2 (iSH2) domain now clashes with the membrane^26^. This interaction between p85α and p110α repress basal PI3K activity but also stabilises p110α, thereby protecting it from degradation, while ultimately being required for the recruitment of p110α to the membrane^36^. Furthermore, the E542K and E545K oncogenic mutations located at this interface relieve p85α(nSH2)-dependent inhibition of p110α^37^. Exposed hits on the helical domain of p110α, [L540P, S541P] and [E547G, K548E/R/G], were validated in a co-competition assay (Figure 4f). These edits, which were confirmed by Sanger sequencing (Figure 4g), lead to downregulation of the protein (Figure 4h) in the absence of changes to mRNA expression (Supplementary Figure 8g). This suggests that the edits lead to post-transcriptional reduction in protein compared to cells edited with a guide against a non-essential gene. Similarly, exposed hits on the C2 domain, [H362R] and [I406T, C407R], were also validated by co-competition assays (Figure 4f) and engendered downregulation of p110α protein (Figure 4h), but not *PIK3CA* mRNA (Supplementary Figure 8g). Another neighbouring patch in p110α showed a significant OR in the helical domain around buried helices: [Y557H, C558R], [C558R], [C558R, V559A], [L565P], [L586S/P, V587A] and [V616A, C618R]. Impact of mutating V616 and C618 was validated (Figure 4f, 4h and Supplementary Figure 8g). The active site mutations [Y836C], [H917R] and [N918S/G/D] (the latter on the catalytic loop) also reduced stability (Supplementary Figure 8e). Our data are consistent with induction of p110α degradation as an alternate therapeutic strategy. In this regard, we note that the orthosteric PI3K inhibitors, inavolisib and taselisib, also induce p110α degradation in a subset of cancer cell lines^38^.

Finally, using the AlphaFold database^39^ structure of MAPKAP1/SIN1 (available PDB structures do not cover the whole protein), which is a critical component of the mTORC2 complex^40^, we identified one OR site in the CRIM (Conserved Region In the Middle) domain (Supplementary Figure 7e and 7g). The CRIM domain of MAPKAP1/SIN1 is predicted to interact with mTOR, which is consistent with the observed low-resolution density. In our case, most hits were located on the central helix: [L206P, I207T] (3 guides), [I210T, C211R], [Q213R, Y214C, T215A], or around the central helix (e.g., [Y230H, C231R]). This region corresponds to a ubiquitin-like fold with a protruding acidic loop, which is considered essential for mTORC2 binding^41^ and in the recruitment of substrates like AKT^42^.

Overall, the combination of CRISPR base editing with the OR patches approach identified both PPI and protein–substrate interactions between mTOR and RHEB, p110α and p85α, and in the MAPKAP1 CRIM domain, suggesting that this workflow can be used to identify important structural interaction sites that may be suitable for drug discovery.

### Patches at intra-protein domain interfaces with unknown biological significance

In p110α, we identified two OR sites of unknown biological significance. The first (p110α site 4) is located between the kinase domain, the helical domain and the Ras binding domain (RBD) (Figure 5a), and next to p110α site 1. The exposed hits in the deep cavity of this region are [F666L/S/P, F667L] (Figure 5b and Supplementary Figure 8g), and [Y270C, K271E/R/G]. Directly at the opening of this pocket, at the back of the C-lobe, there are hits that are cross-detected in p110α site 1: [W880R] (Figure 5b–d and Supplementary Figure 8g), [K884R, N885D/G/S, K886E] (Figure 5b–d), [N885D/G/S, K886E/G/R] and [D926N, G927E/K/R] (Figure 5b, 5d and Supplementary Figure 8g). The affected residues are spatially close to the catalytic domain and possibly affect its shape. We believe we are the first to describe a biologically relevant impact to them. Edits from all sgRNAs impacted p110α activity and protein stability post-transcriptionally (Figure 5d, Supplementary Figure 8g).

**Figure 5:**
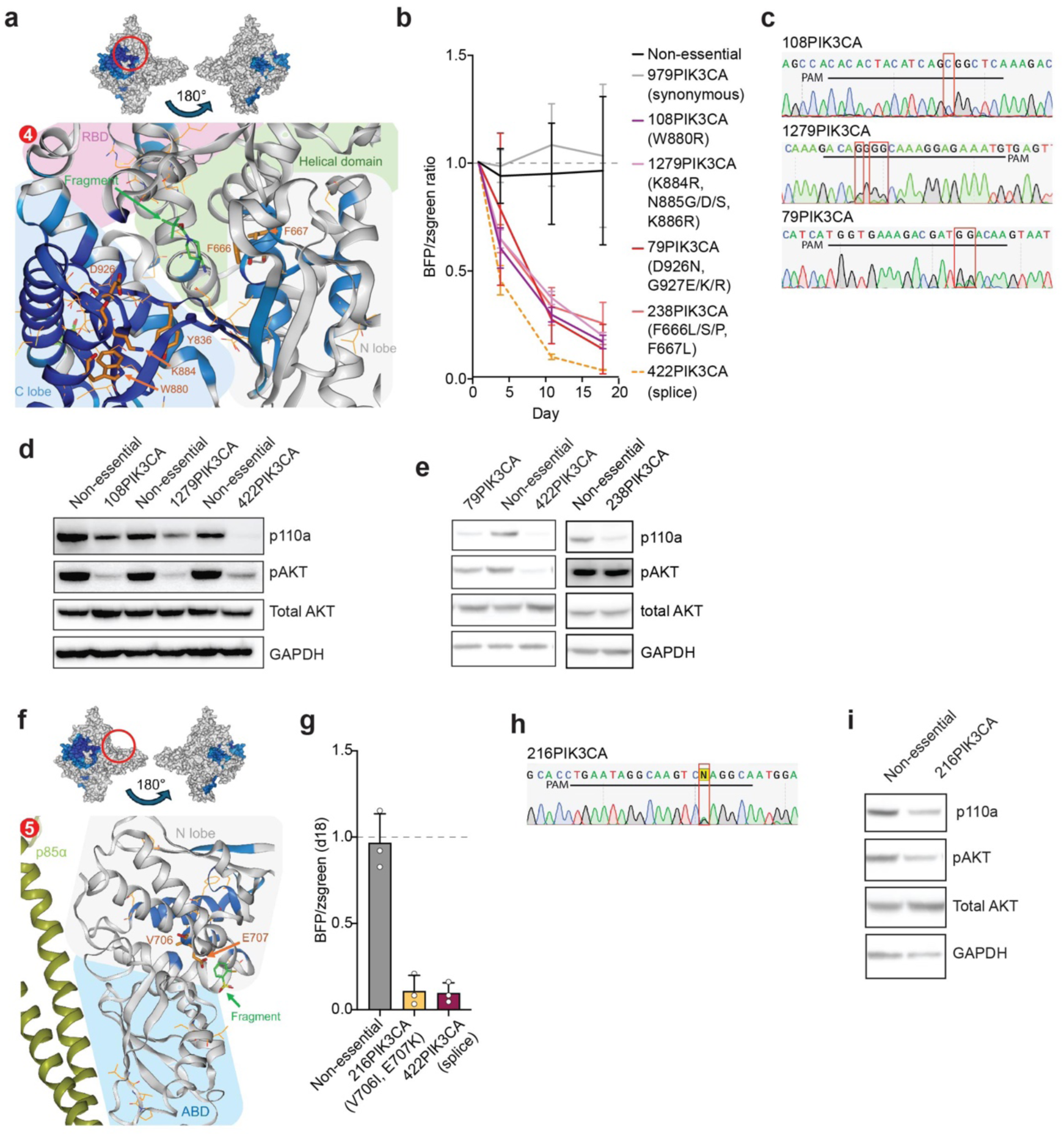
Structural analysis on two additional fragment sites of p110α. **a**) A miniature surface representation of p110α is shown to highlight the location on top and a cartoon representation is coloured by OR below. Shown is the fragment site behind the kinase site between the helical domain and RBD domain. **b**) Co-competition experiment displays the proportion of MCF7 ABE/CBE cells transfected with tested sgRNA relative to non-essential sgRNA. N=2 for 108PIK3CA, 1279PIK3CA, 79PIK3CA (CBE), 238PIK3CA, 979PIK3CA (synonymous control) and 422PIK3CA (splice control). **c**) Sanger sequencing of MCF7 ABE/CBE cells transfected with tested sgRNAs. **d&e**) Western blot of MCF7 ABE or HGC27 CBE cells transfected with tested sgRNAs 7 days after transfection. **f**) A miniature surface representation of p110α is shown to highlight the location on top and a cartoon representation coloured by OR is below. Shown is the fragment site between the N-lobe and ABD. **g**) Co-competition experiment displays the proportion of MCF7 CBE cells transfected with tested sgRNA relative to non-essential sgRNA at day 18, N=3. **h**) Sanger sequencing of MCF7 CBE cells transfected with tested sgRNAs. **i**) Western blot of HGC27 CBE cells transfected with tested sgRNA 7 days after transfection.

The second p110α OR site of previously unknown significance, p110α site 5, is located between the N-lobe (next to the helical domain) and the adaptor-binding domain (ABD) (Figure 5f). A guide hit in this site located at the edge of the significant OR patch is [V706I, E707K] (Figure 5f and Supplementary Figure 8g). E707 is an exposed residue and part of an allosteric pocket proposed by Zhang et al.^43^ that has potential for ligand interactions. Recently a key salt bridge between C2 domain R349 and kinase domain E707 has been proposed to be central to PI3Kα homodimer formation and PI3Kα activity^44^. The effect of V706 and E707 mutations on endogenous PI3Kα signalling and cell proliferation has not previously been shown. The impact of the edits from all validated guides on p110α activity and stability was at the post-transcriptional level (Figure 5g and Supplementary Figure 8g).

Whilst the biological impact of perturbing both regions was previously unknown, both sites have hits bound from a fragment screen reported by Miller et al.^45^; p110α site 4 (see PDB 5SWP, visible in Figure 5a) and site 5 (PDB 5SW8, visible in Figure 5e). We propose that combining data from CRISPR and fragment screens could be used to help prioritize fragments that hit biologically relevant regions and are therefore good starting points for a drug discovery campaign.

We also found two OR patches of previously unknown biological significance close to PPIs within mTOR. mTOR site 3 is at the intersection between the FAT domain (that precedes the N-lobe region in the amino acid sequence) and the C-lobe, close to the mTOR stabilising InsP6 site^46^ and not far from the DEP1 domain of the mTOR-inhibitory protein DEPTOR (Figure 6a). Many of the hits within the OR patch are buried within the protein, that is [T1903I, L1904F], [L1904P], [L1907F], [W1935R], [L1956P, I1957T] and [L1960P], and are likely to impact protein stability, especially given the frequency of leucine to proline mutations. Close to this OR patch, we identified the surface exposed hits [R1905G] and [D1902G, T1903A] (Figure 6b−d). Mutations of both surface exposed residues led to reduction of mTOR activity with minimal impact on mTOR stability (Figure 6d). mTOR site 4 lies at the intersection between the FAT and M-HEAT domains, opposite the RHEB interface site (Figure 6e), and adjacent to a deep pocket: [L1317P, F1318S/L/P] and [L1317P] (Figure 6f and 6g). F1318 is located in a hydrophobic region, but it is a partly exposed and could therefore be targeted with π-stacking or lipophilic interactions. From western blotting, we observed that both mutations affected mTOR activity and had a minor impact on protein levels (Figure 6g).

**Figure 6:**
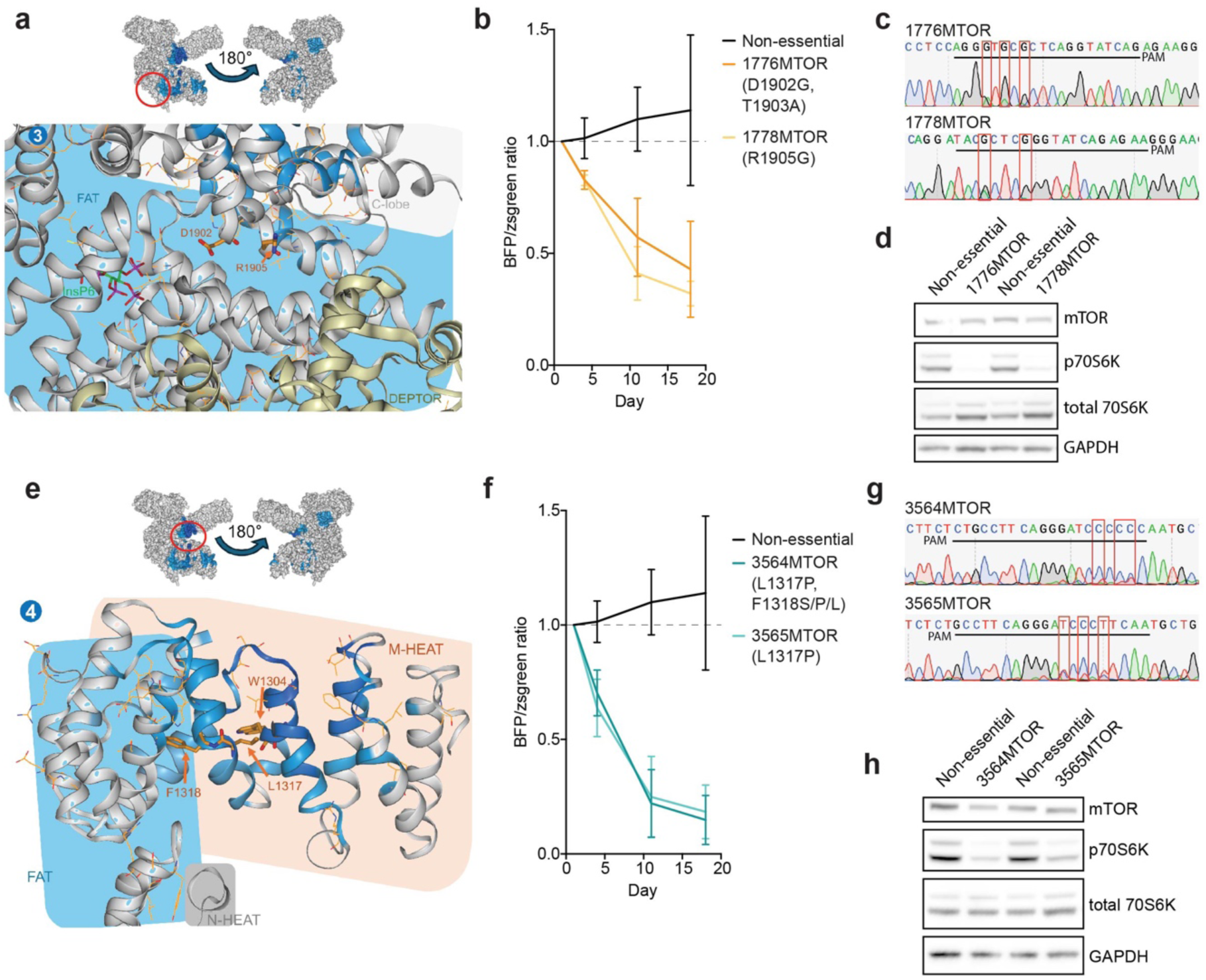
Structural analysis of screening results on new sites. **a**) A miniature surface representation (top) and cartoon representation (bottom) of a new mTOR site between the kinase C-lobe and FAT (PDB code: 7PE9). **b**) Co-competition experiment displays relative growth of target sgRNAs compared with non-essential sgRNA, N=3. **c**) Sanger sequencing of edited MCF7 ABE cells. **d**) Western blot of edited MCF7 ABE cells 7 days after transfection. **e**) A miniature surface representation (top) and cartoon representation (bottom) of a new mTOR site between the N-HEAT, FAT and M-HEAT domains behind the mTOR–RHEB interface (PDB code: 6BCU). **f**) Co-competition experiment displays relative growth of target sgRNAs compared with non-essential sgRNAs, N=3. **g**) Sanger sequencing of edited MCF7 ABE cells. **h**) Western blot of edited MCF7 ABE cells 7 days after transfection.

In PDPK1, one significant patch was identified when using the 3QD0 PDB structure (Supplementary Figure 9e–g). Hits within the patch include [K115E/G/R], [E121G, N122D/G/S], [R129K, E130K], [E130G, R131W], and [E130K, R131Q]. These hits mutate charged amino acids (explicitly annotated as “polar residues” in Uniprot) to other amino acids with identical or opposite charges. The interaction between AKT1 and other proteins with pleckstrin homology (PH) domains (such as PDPK1) relies on hotspots of charged interactions^47^. We were unable to retrieve the same significant patch when using the 3NAX PDB structure, that displays a different conformation of a loop in the N-lobe, reducing the spatial clustering of the identified hits. Notably, in 3QD0, the E130 residue in the N-lobe is bound to the inhibitor and is present in a helical conformation, which faces inwards. In contrast, E130 is in a more disordered loop conformation and oriented towards the surface in the 3NAX structure. The flexibility of this loop might explain its ability to both interact with ligands and form a PPI with the PH domain. We also observed a few hits that do not form part of the OR patch (Supplementary Figure 5g, 5h, 7d and 7f) in the region of the protein which may interact with the PH domain^47^. This highlights how the structure used can impact the identification of OR patches and the importance of using multiple structures if available.

Overall, we identified and validated OR patches in regions of proteins that have not previously been ascribed a biological function. These regions offer potential novel points for intervention with a small molecule approach.

## Discussion

Understanding how specific amino acid residues contribute to protein function remains a central challenge in translating genomic knowledge into therapeutic opportunities. Despite important progress in computational approaches such as AlphaFold^48^ and AlphaMissense^49^, predicting functionally impactful residues is difficult because protein activity is shaped by long-range molecular interactions that influence 3D structure, intra-molecular and multi-protein interactions, and due to the dynamic nature of proteins themselves. In this study, we combined pooled CRISPR base editing screens with structural analysis to systematically map functional residues across the class IA PI3K signalling pathway in multiple disease relevant models. We introduced thousands of predicted missense mutations across all 30 genes of the pathway at semi-saturating coverage. Notably, this allowed us to interrogate the impact of mutations on the endogenous protein at scale, without the need for protein overexpression. This approach preserves multi-protein complex stoichiometry and the *in cellulo* dynamic states of proteins, which can be challenging to maintain using conventional biochemical and biophysical approaches, and therefore has the potential to reveal protein structure-function insights that would otherwise be inaccessible.

In addition to providing a resource of individual hit residues, mapping functional variants onto protein structures enabled identification of spatially clustered “hotspots” enriched for deleterious mutations (referred to as OR patches). Compared to a previously published 3D hierarchical clustering method (Ngan et al^50^), our less computationally heavy and easily interpretable approach allowed us to perform structural mapping at scale and to identify clusters of pocket-like size with a significantly higher fraction of hits, providing a narrower follow-up search space for potential drug discovery approaches^51,52^. Focusing on small phenotypically relevant regions can reduce the computational load for downstream approaches such as high-throughput virtual screening. Our approach can be applied to many types of structures including NMR, cryo-EM, X-ray crystallography or AlphaFold predicted models. We noted that the exact model used can impact OR patch retrieval, as was seen for PDKP1, highlighting the importance of comparing multiple structures if possible.

The most prominent clusters occurred within kinase domains, consistent with their central role in enzymatic activity. Whilst orthosteric inhibitors are valuable, allosteric compounds are often more selective, as exemplified by MEK1/2 inhibitors trametinib and selumetinib^53^, and can help combat drug resistance as exemplified by BCR-ABL inhibitor asciminib^54^. We identified multiple enriched regions outside of catalytic pockets, with several located at intramolecular domain junctions. In p110α an allosteric pocket has been predicted between the N lobe and ABD, but functional relevance has not been shown^43^. We identified an OR patch covering part of this pocket and showed that mutation of residues V706 and E707 within this patch caused reduced p110α activity. Another OR within p110α had multiple exposed hits that are spatially close to the catalytic site and hence may impact its shape. Edits within both regions impacted protein stability post-transcriptionally. Compounds that impact protein stability and degradation are a growing area of research and several allosteric inhibitors have been identified, such as recent covalent and non-covalent WRN inhibitors^55,56^.

In our data, we also identified clusters at PPI interfaces. Targeting PPIs can be an effective strategy for modulating pathway activity, as demonstrated by BCL-2 family mimetics such as navitoclax and venetoclax, and tri-partite RAS inhibitors. Such regions may provide opportunities to target specific multiple protein complexes which may share common kinases and elicit different phenotypic responses when targeted. This is exemplified by mTORC1 and mTORC2 complexes, which share the common kinase mTOR, but have specific signalling functions^57^. We identified an OR patch within the mTOR-RHEB interface. Targeting this interface has the potential to specifically target mTORC1 but has historically been challenging due to the large interaction surface, though one small molecule and a peptide have been identified^33,58^. We also found other OR hits in mTOR that may impact protein—protein interfaces. mTOR site 3 is close to both the InsP6 binding site within mTOR and the DEP1 domain of DEPTOR. First, due to the proximity of D1902 to the InsP6 interaction site, disruption of this might disturb the mTOR-stabilising InsP6 interaction^46^. InsP6 interacts with multiple lysine and arginine residues in mTOR including K1788 which was a hit in the screen ([K1788G]). Secondly, molecular dynamic simulations of mTOR indicate a potential salt bridge between R1905 of the FAT domain and E2419 of the kinase domain. Mutation to E2419K is predicted to result in a weaker interaction between FAT and N-lobe and predicted to increase mTOR activity^59^. However, we observe the opposite impact on mTOR activity when mutating R1905 *in vitro*. This may be due to impacts on the orientation of catalytic residues or targeting the pocket towards the R1905 could alter the interaction with inhibitory protein DEPTOR^60^. In mTOR site 4, F1318 is partly buried under the flexible R1315 on the M-HEAT domain that interacts with FAT residues E1352 (visible in PDB structure 7PE9) or T1356 (visible in PDB structure 6BCU). Targeting this potential site is likely to alter mTOR tertiary structure and prevent mobility of the FAT domain relative to the M-HEAT domain, which is a key determinant of mTOR activation following RHEB binding^31^.

Our results also illustrate how functional genomic screens can complement other drug discovery approaches. Several OR patches overlapped with fragment-binding sites identified in previous structural studies, suggesting that integrating cellular perturbation data with structural and chemical datasets may help prioritise ligandable regions most likely to influence protein function. This concept is particularly relevant for multidomain proteins such as mTOR or PI3K, in which structural pockets are abundant but only a subset of which are functionally critical, and for proteins that lack precedent for druggability, such as transcription factors. More broadly, our approach is highly complementary to Fragment-Based Drug Discovery (FBDD), high-throughput screening (HTS), DNA-encoded library (DEL) screening and virtual screening, by helping to triage and to nominate candidate drug-binding sites and guide medicinal chemistry efforts. We anticipate that integrating functional genomics with structural and chemical biology will help prioritise druggable, functionally relevant sites and accelerate the development of new therapeutics, particularly for currently difficult to drug targets.

Despite these advances, several limitations remain. Base editing is constrained by the spectrum of accessible mutations and can introduce bystander edits, resulting in incomplete or ambiguous coverage of some residues. Ongoing improvements in genome-editing technologies, including expanded PAM recognition, prime editing and higher efficiency editors, should enable more comprehensive mutational coverage^61,62^. In addition, the OR framework used to detect spatial clusters prioritises regions enriched for multiple hits and may overlook individual functionally critical residues. This limitation may be particularly relevant for residues such as cysteines, which are valuable in covalent drug discovery^63^. Independent experimental validation of variant effects remains a major bottleneck. Scalable, systematic validation approaches using orthogonal functional readouts, such as measurements of protein stability and effector binding^64^, could further enhance and refine interpretation of mutational effects.

In conclusion, our proof-of-concept study suggests a future in which large-scale perturbation datasets in cells are combined with structural modelling, *in silico* prediction and chemical biology to generate predictive maps of protein function that accelerate the discovery of new therapeutic strategies targeting disease-linked signalling networks in oncology and beyond.

## Methods

### Cell culture

MCF7 and T47D (NCI) cells were cultured in RPMI 1640 (ThermoFisher, 52400025) media supplemented with 10% FCS (ThermoFisher, A5256701), 1% penicillin/streptomycin (ThermoFisher, 15140122), 4.5 mg/ml glucose (Sigma Aldrich, G7528-250G) and 1 mM sodium pyruvate (ThermoFisher, 11360039). HGC27 (RIKEN) cells were cultured in Dulbecco’s Modified Eagle’s Medium (DMEM/F12, ThermoFisher, 31330038) supplemented with 10% FCS and 1% penicillin/streptomycin. All lines were verified by STR profiling and mycoplasma tested. Selection with blasticidin (InvivoGen, ant-bl-1), puromycin (InvivoGen, ant-pr-1) or hygromycin (Toku-E, H011-20ml) was performed using the doses indicated in the relevant Methods subsections.

### Base editing cell line generation

Base editing cell lines were generated as previously described^65^. Briefly, prior to transfection, cells were treated overnight with 1 µM AZD7648 (MedChemExpress, HY-111783) to increase homologous recombination rates. Plasmids encoding Cas9 and the human CLYBL locus sgRNA (5′-ATGTTGGAAGGATGAGGAAA-3′) and plasmids encoding tet-ON base editor (BE3.9max-Cas9NGN^14^ or ABE8e-Cas9NGN^15^) within CLYBL homology arms were co-transfected using FugeneHD (Promega, E2311). Transfected cells were selected with blasticidin (10 µg/ml MCF7, 10µg/ml T47D and 20 µg/ml HGC27) for four days and then maintained in media with half the concentration of blasticidin used for selection. mApple-positive cells were selected by FACS. Clonal lines were generated and editing efficiency was assessed using BE-FLARE^16^ (CBE) or stop codon GFP reporter^17^ (ABE). Plasmids used in this study can be found in Supplementary Table 3.

### CRISPR base editing library production

CRISPR sgRNAs were designed using BEstimate^19^ (https://github.com/CansuDincer/BEstimate) against the human reference genome GRCh38; we assumed a wild-type genome and a predicted editing window of 4 to 9 nucleotides. NGN guides were used for more flexible sgRNA design across the targeted genes. sgRNAs were then designed and filtered to extract those that target the coding sequence (CDS). To generate control essential splice and non-essential splice sgRNAs, SpliceR^66^ was used and the top three sgRNAs were ranked using the following metric [cDNA disruption score x (ABEscore + CBEscore)]. Oligo pools (Twist Biosciences) were PCR amplified (KAPA HiFi HotStart ReadyMix, Roche) and inserted into a Bbs-I digested pKLV2-BFP-T2A-puroR lentiviral backbone (Addgene #67974) using Gibson assembly (NEB). After ethanol precipitation, multiple electroporations (Endura Competent Cells, Lucigen) were performed and plasmid pools were propagated in LB with 100 µg/ml ampicillin at 32°C overnight and extracted (Qiagen Plasmid Maxi Kit).

For virus packaging, HEK293T were co-transfected with the sgRNA plasmid pool, psPAX2 and pMD.2G using FuGeneHD (Promega) in Opti-MEM (Thermo Fischer Scientific). The media was changed the next day, and viral supernatant collected 72 h post-transfection.

### CRISPR base editing screens

All screens were performed at approximately 1000x coverage. Cells were transduced with 8 μg/ml polybrene (SantaCruz, 28728-55-4) and a viral sgRNA titre that achieved 30−50% transfection rate, as measured by BFP fluorescence 48h post-transduction. 48h post-transduction, cells were selected with puromycin for 4 days and, at the same time, base editing was induced with 1 µg/ml doxycycline (Toku-E, 24390-14-5) for 4 days, with the exception of MCF7 CBE and T47D CBE which were induced for 7 days. Each screen was independently repeated twice. Cell pellets were collected at day 17 post-transduction for HGC27 and day 24 for MCF7 and T47D.

DNA was extracted (DNeasy Blood and Tissue, Qiagen, 69504), sgRNA sequence PCR amplified (PCR1, 28 cycles, 3 µg per reaction, Q5 Hot Start High-Fidelity, M0494S) with multiple reactions to maintain coverage. Plasmid DNA from the original library served as a control in the screening experiments. Column purified (Qiagen, 28104) PCR1 products were then indexed with a second round of PCR (8 cycles, 1ng per reaction, KAPA HiFi HotStart ReadyMix, Roche), SPRI purified (AMPure XP SPRI beads; Beckman Coulter) and quantified (Bioanalyzer; Agilent). Libraries were sequenced at 1000 to 2000x coverage on HiSeq2500 (Illumina) using 19 bp single end reads sequencing on Rapid Run Mode with a custom sequencing primer (5 - TCTTCCGATCTCTTGTGGAAAGGACGAAACACCG-3).

### CRISPR base editing screen analysis

MAGeCK analysis^67^ was performed using default parameters, except normalisation was set to ‘none’, as the input corrected counts had already been normalised. Read counts were normalised by generating reads per million (RPM) using reads/total reads for sample x (1,000,000 + 1 pseudo count). Average log_2_ fold-change (LFC) values as [log2(RPM condition / RPM plasmid control)] were then calculated. For replica screens, the average LFC was taken. Z-scores were calculated as [(LFC - mean LFC) / sd], where sd is the standard deviation of the LFC values for that screen. Any guides with fewer than 50 counts in the control plasmid library were excluded.

The fasta and gff3 files for the GRCh38 genome assembly were downloaded from the ftp server of the Ensembl release 115. Off-targets were calculated with Casofffinder^68^ v2.4.1, mapping location on the GRCh38 chromosomal and mitochondrial DNA using the gff3 files. Any guide with zero or one mismatch against the coding region of any essential genes for each cell line (Supplementary Table 4) was filtered out, as were guides with four or more off-targets with zero or one mismatch. This led to removal of 822 of guides, 2% of the library (with a few exceptions, such as RPS6 or EIF4E having 55% and 25% of off-target guides, respectively).

Post-screening, we noted many internal positive controls against essential genes failed to give a signal. Data across multiple cell lines^55,69^ was compiled and the top 150 splice essential guides for ABE or CBE was taken. Guide sets were then intersected, and those found in both ABE and CBE were taken forward. This led to a sub-selection of 110 splice essential positive control guides (Supplementary Table 2). Based on the poor discrimination between negative and positive controls observed in CBE for the T47D cell line, we decided to remove this combination from the downstream analysis.

To assign hits, a LFC cutoff defined as a False Discovery Rate (FDR) of 5% (i.e. the highest LFC at which the negative controls represent up to 5% of all controls) was calculated for each screen using all negative control guides and the sub-selection of positive control guides, similar to that which was performed by Cuella-Martin et al.^70^. This corresponded to an LFC of -2.02 and -0.429 for HGC27 ABE and CBE, respectively, -1.53 and -0.51 for MCF7 ABE and CBE, respectively, and -0.95 for T47D ABE. Guides with an LFC equal to or below this value and a significance of p < 0.05 were called as hits.

The impact of each mutation was assessed using Ensembl Variant Effect Predictor (VEP)^22^ with the bioconda package ensembl-vep 115.2. When calling the potential impact of a guide, the most severe consequence was selected with the following order: splice/stop > missense > synonymous.

The essentiality for each gene was determined from DepMap data^21^, where all genes with a gene effect lower than -0.5 were considered essential.

As the predicted outcomes of sgRNAs can vary dependant on if all or some bases are edited within the base-editing window, unless validated, the nomenclature within the manuscript used to refer to mutations generated per sgRNA will be [mutation 1, mutation n], where each mutation is defined by wild-type amino acid, residue number, and potential mutated amino acids introduced.

### Structural analysis

For each guide–editor combination in the screening, we computed the percentage of hit cell lines in ABE (e.g. 0−3 cell line hits corresponding to 0, 33, 66, or 100%) and in CBE (e.g. 0−2 cell lines hits corresponding to 0, 50 or 100%). For each mutated amino acid position induced by missense edits, the maximum percentage of hit cell lines among all corresponding guides was associated (e.g. if an amino acid is mutated by two guides, where a guide hits 66% of cell lines in ABE while the other guide hits 50% of cell lines in CBE, 66% is associated to that amino acid position). For each screened gene, plots were generated to visualise the minimum (across all guides) Z-score per residue for all guides for each cell line in CBE and ABE, along with Uniprot annotations. See Supplementary table 5 for Protein Data Bank (PDB) and AlphaFold database codes used.

To identify areas of the proteins that were enriched in hits, we scanned each full protein structure using a patch approach: centring on each individual residue, neighbouring residues having an atom within 7 Angstrom of any central residue atom were compiled into the patch. A Fisher statistical test was conducted to compute the odds ratio (OR, with corresponding p-values) as follows:

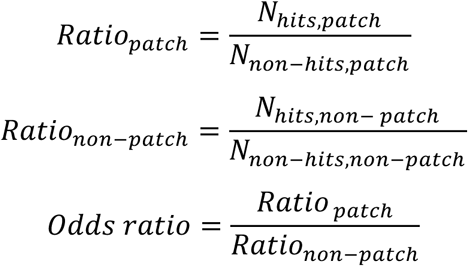

For a full protein structure, the p-values were corrected using the Benjamini-Hochberg method with an alpha of 0.05 using the statsmodels v0.14.2 Python package, and patches with corrected p-values < 0.05 were identified and mapped into the structures (i.e., the fraction of hits that is significantly higher in the patch compared with the rest of the protein).

A similar approach was used by Ngan et al., who performed 3D hierarchical clustering of guide RNAs in a CRISPR tiling assay, using structural information, by weighting the Euclidean distance between residues with pairwise guide RNA scores ^50^. Our approach is simpler to compute and interpret but is sensitive to the LFC threshold (to define hits vs non-hits) and the radius of the patch. While their hierarchical approach can separate subpockets within pockets and identify smaller local hit cluster without any degree of significance, our approach iterates over patches of residues of a given size and identify clusters having a higher concentration of hits with associated p-value.

### Recombinant sgRNA plasmid cloning for validation experiments

Individual sgRNAs were cloned in an arrayed format using Golden Gate assembly into the relevant pKLV2-hU6-sgRNA expression vector. Primers encoding the sgRNA and BbsI overhangs (Forward: 5 -CACCGNNNNNNNNNNNNNNNNNNN-3 and Reverse: 5 -AAACNNNNNNNNNNNNNNNNNNC-3) were annealed and ligated into BbsI entry vectors using Golden Gate cloning (BbsI-HF and T4 DNA ligase from NEB in 30 restriction and ligation cycles). Reactions were transformed in DH5-alpha *E*. *coli* (NEB, C2987H) and clones were sequence verified (Eurofins Genomics). Recombinant vector and oligonucleotide sequences can be found in Supplementary Table 3 and 6.

### Lentivirus production for validation experiments

HEK cells were plated in 12-well plates and transfected with a mixture of 375 ng lentiviral construct containing sgRNA, 455 ng psPAX2 (Addgene #12260), 75 ng pMD2.G (Addgene #12259) and 2 µl FugeneHD (Promega, E2311) in 30µl OptiMEM (ThermoFisher, 15392402). The supernatant was filtered through a 0.2µm filter after 2 days incubation and frozen at -80°C.

### Co-competition assay

MCF7 ABE or MCF7 CBE cells were transduced at a high multiplicity of infection (MOI) with lentivirus of individual sgRNAs. After 24 h, cells were selected with puromycin or hygromycin and base editing was induced with 1 µg/ml doxycycline for 4 days. Cells were mixed day 5 post-transduction and ratios of blue fluorescent protein (BFP) to zsgreen or mAzami quantified on day 6 post-transduction. The ratio of BFP to zsgreen/mAzami was tracked over time using flow cytometry.

### Western blotting

MCF7 ABE or HGC27 CBE cells were transduced at a high MOI with lentivirus of individual sgRNAs, selected with puromycin or hygromycin after 24 h and base editing was induced with 1 µg/ml doxycycline. Non-essential sgRNAs were used as a control. Cells were lysed at D7 in 1X RIPA buffer supplemented with 2x Halt protease and phosphatase inhibitor (ThermoFisher, #78440). Lysates were prepared with 4X loading buffer and 10% beta-mercaptoethanol or 1X NuPAGE reducing agent and run on NuPAGE 4-12% Bis-Tis gels (ThermoFisher, NP0322BOX). PVDF membranes were blocked in 5% milk in TBS-T and probed with the following primary antibodies: p110α/*PIK3CA* (CST, C73F8, 1:1000), AKT (CST, 9272S, 1:1000), pAKT-S473 (CST, 9271S, 1:1000), RHEB (CST, 13879S, 1:750), MTOR (CST, X, 1:1000), p70S6K-T389 (CST, 9234S, 1:1000), p70S6K (CST, 9202S, 1:1000). GAPDH (CST, 5174S, 1:2000) was used as a loading control. The membranes were washed with TBS-T and incubated with 1:2500 anti-rabbit IgG HRP-linked secondary antibody (GE Healthcare, NA931V-ECL HRP) in TBS-T + 5% milk. Signal was visualised using a Cytive Amersham ImageQuant system.

### qPCR

MCF7 ABE or HGC27 CBE cells were transduced at a high MOI with lentivirus of individual sgRNAs, selected with puromycin or hygromycin after 24 h and base editing was induced with 1 µg/ml doxycycline. RNA was extracted from snap frozen cells (Qiagen 74134), quantified and cDNA generated from 1µg of RNA (Qiagen 205311). 1 µl of 1:10 diluted cDNA was used with TaqMan master mix (ThermoFisher, 4444556) and probes (*PIK3CA* TaqMan probe, Applied Biosciences, Hs00907957_m1; GAPDH TaqMan probe, Applied Biosciences, Hs00266705_g1; PUM1 TaqMan probe, Applied Biosciences, Hs00472881_m1) according to the manufacturer’s protocol in a 96-well format (ThermoFisher, 4346907). The qPCR was run using standard protocol on a Quantstudio5 (ThermoFisher).

### Sanger sequencing of edited sequences

Cells were transduced at a high MOI with lentivirus of individual sgRNAs, selected with puromycin or hygromycin after 24 h and base editing was induced with 1 µg/ml doxycycline. DNA was extracted (Qiagen, 69504) and used for PCR amplification of the edited region (oligonucleotide sequences in Supplementary Table 6) by Q5 high-fidelity polymerase (NEB, M0492S). PCR products are purified (Qiagen, 28104) and sent for Sanger sequencing (Eurofins Genomics). Sequence traces were analysed using SnapGene® software (from Dotmatics; available at snapgene.com)

### Software

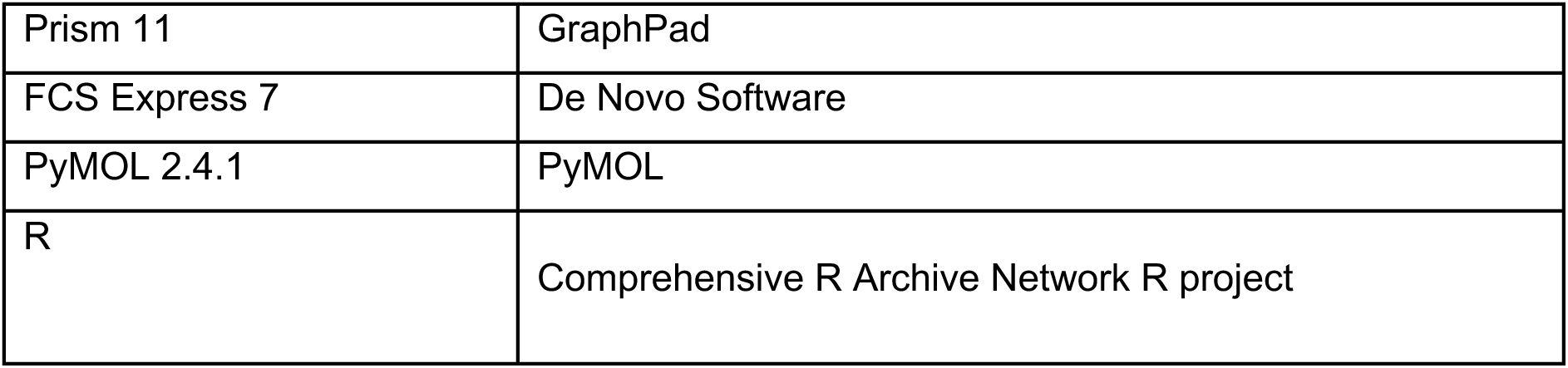

|  |  |
| --- | --- |
| Prism 11 | GraphPad |
| FCS Express 7 | De Novo Software |
| PyMOL 2.4.1 | PyMOL |
| R | Comprehensive R Archive Network R project |

## Data and code availability

Sequencing data from base editing screens will be made available (ENA accession numbers in Supplementary Table 7) and analysed datasets are available in Supplementary Table 8. The base editing screen and structure analysis code is available on GitHub. https://github.com/Garnett-Lab/crispr-be. The authors have applied a AGPLv3 license to all code in this study.

## Acknowledgements

K.M. and B.A. were Sustaining Innovation Postdoctoral Research Associates at Astex Pharmaceuticals and thank Astex Pharmaceuticals for funding. We thank the Garnett laboratory and wish to acknowledge the contribution of the Cancer Ageing and Somatic Mutation Support team at the Wellcome Sanger Institute. We thank the Sanger Flow Cytometry Facility and the Sanger Core Sequencing pipeline. This work was funded in part by Wellcome Trust Grant 206194.

## Author contributions

KM, BA, BB and M.J.G devised the study. KM: performed and designed base editing screens and initial data preprocessing. BA: designed the computational analysis procedure, data processing and analysis. BB: Data analysis, image production. KM and LK performed and analysed validation experiments. AW and SW assisted with experiments. C.D., M.A.C, M.S., and G.P. assisted with sgRNA library design and cloning. E.K performed data analysis. BB, KM, LK, BA and M.J.G. wrote, and all authors reviewed, the manuscript. TD, MW, FL, MV: manuscript review.

