## Supplementary_figures for "Structure-function analysis of PI3K signalling cascade base editing screens in cancer cells"

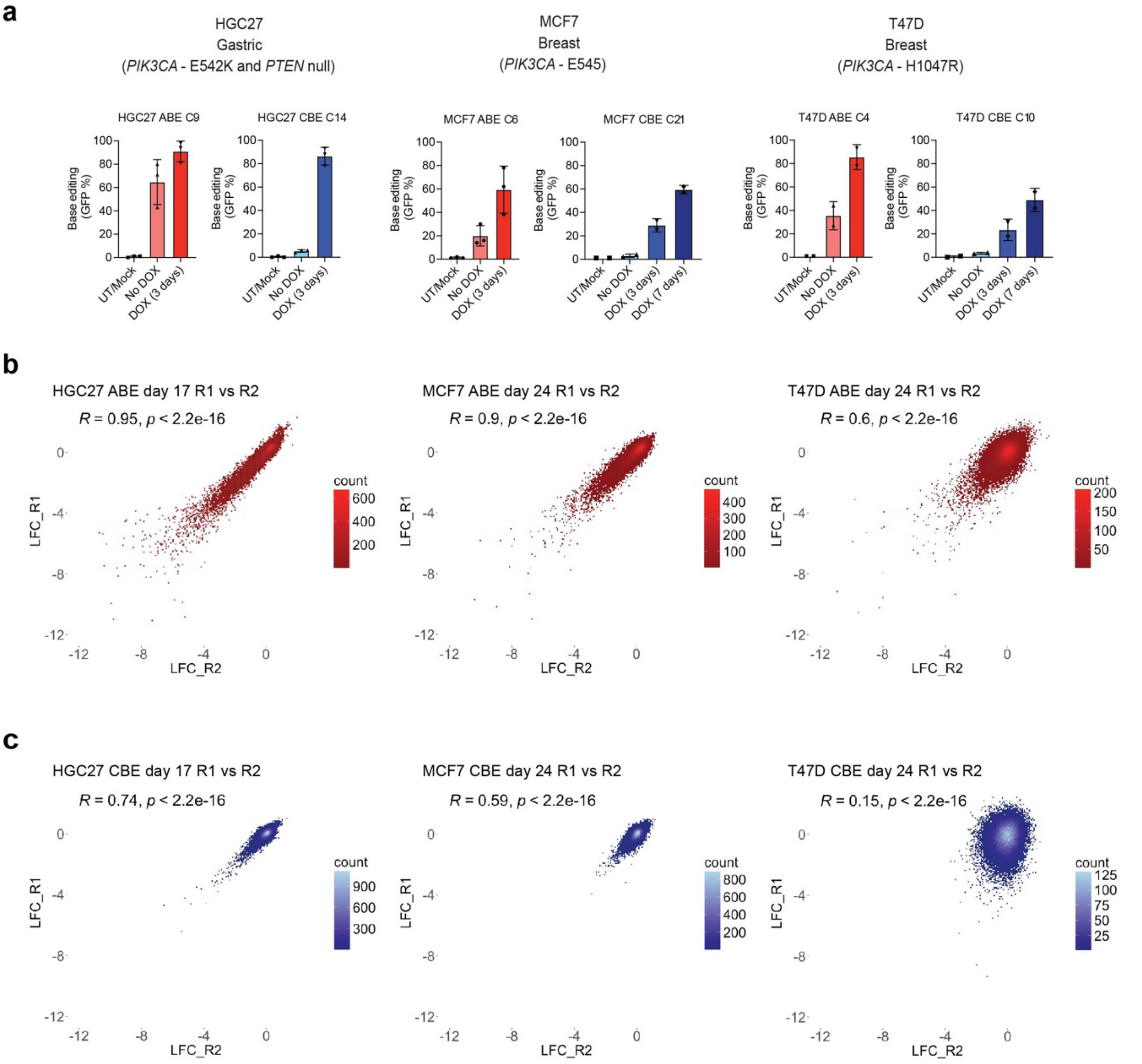


**Supplementary Figure 1: Generation of base editing cell lines and quality control of CRISPR screens. a**) Generation of clonal base editing cell lines from *PIK3CA* mutant cancer cell lines HGC27, MCF7 and T47D. The editing efficiency of individual clones was assessed using BE-FLARE for CBE (blue bars) and GFP stop reporter for ABE (red bars). Readout was performed after 3 or 7 days of doxycycline induction of base editing, with the % of GFP as a read out of editing efficiency. Data are representative of 2 or 3 independent experiments. **b**) and **c**) Pearson correlation of screen replicates comparing plasmid vs end point performed in b) ABE (red) and c) CBE (blue) HGC27, MCF7 and T47D cell lines. Data are representative of two independent experiments. ABE: Adenine base editing; CBE: Cytidine base editing; DOX: doxycycline; GFP: Green fluorescent protein; LFC: Log2 fold change

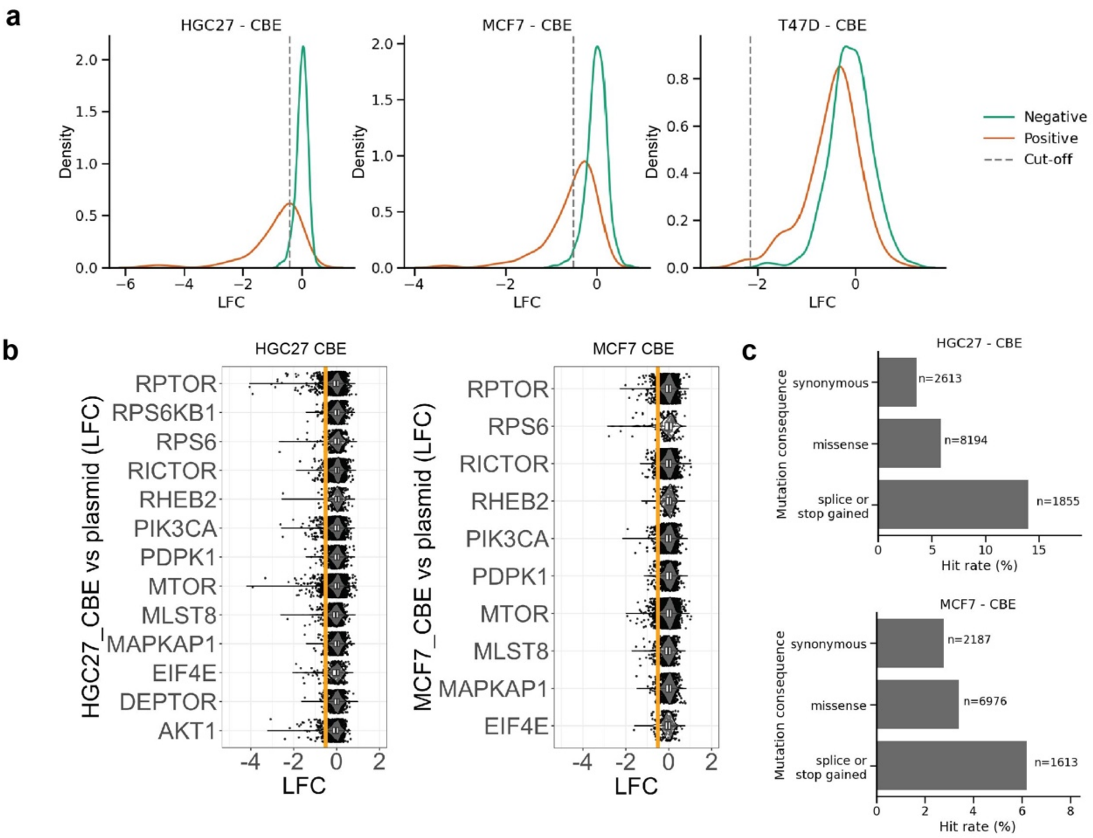


**Supplementary Figure 2: Cytidine base editing screening results on the PI3K signalling pathway**. **a**) Distribution plots of positive and negative screen control guides for CBE screens in HGC27, MCF7 and T47D cell lines. The 312 negative controls include nonessential splice, intergenic and non-targeting sites, whilst the 110 positive controls are essential splice sites. The intersection of these was used to generate an LFC cut off with a 5% FDR for hit calling. **b**) Violin plots showing the depletion of guides across all essential genes within the PI3K pathway for the CBE editor in HGC27 and MCF7 cells. Boxplots show the median and interquartile range. The LFC cut off for hits is an orange line. **c**) Distribution of percentage of hits by most severe predicted edit type for the essential genes highlighted in **b**). The essentiality for each gene was determined from DepMap data[21] using a cutoff of -0.5 to determine essential genes.

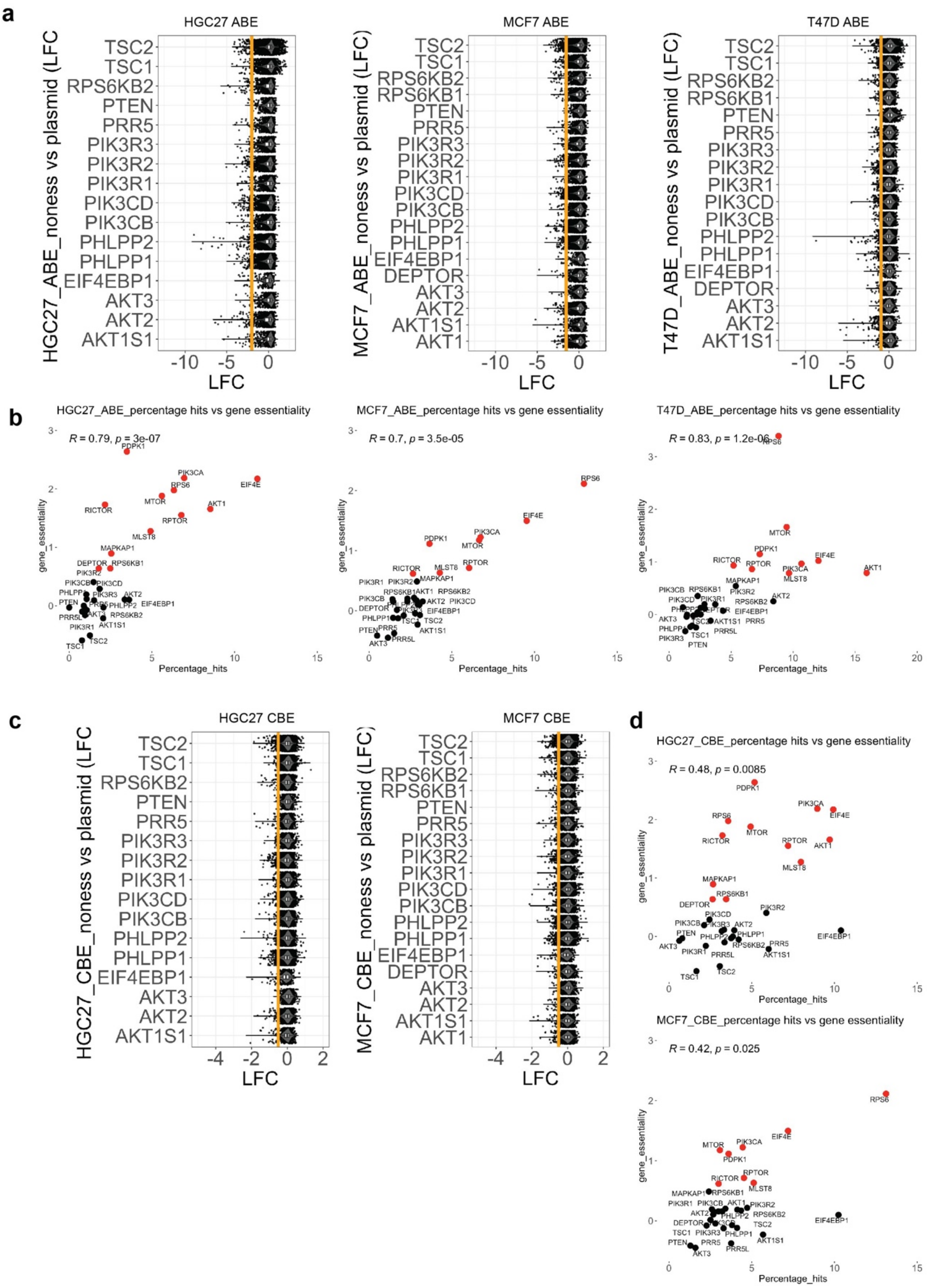


**Supplementary Figure 3: Cytidine base editing screening results on the PI3K signalling pathway (continued). a**) and **c**) Violin plots showing the depletion of guides across all non-essential genes within the PI3K pathway for ABE (a) and CBE (c) editor in HGC27, MCF7 and

T47D (ABE only) cells. Boxplots show the median and interquartile range. LFC cut off for hits is an orange line. **b**) and **d**) Percentage of hit guides for each gene-by-gene essentiality in ABE (b) and CBE (d) screens. Black dots show non-essential genes whilst red dots show essential genes that are components of the PI3K pathway. R is spearman correlation. Gene essentiality is taken from DepMap screening data[21] and essential genes are those with a gene effect of less than -0.5.

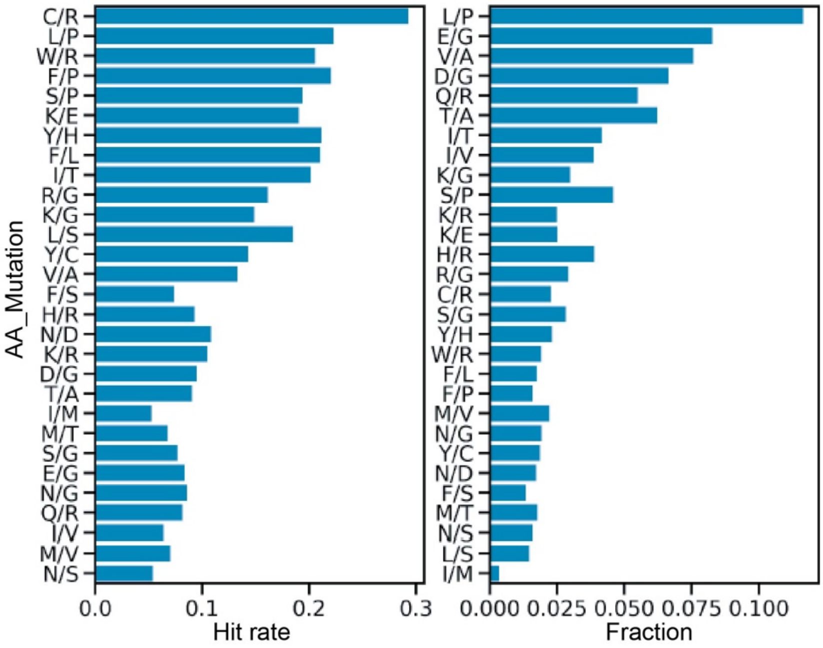


**Supplementary Figure 4:** Hit rate (left panel; fraction of guides that are hit for each corresponding mutation) and fraction of guide inducing given mutations (right panel; proportion of the mutation among all observed mutations) in ABE screens. AA: amino acid

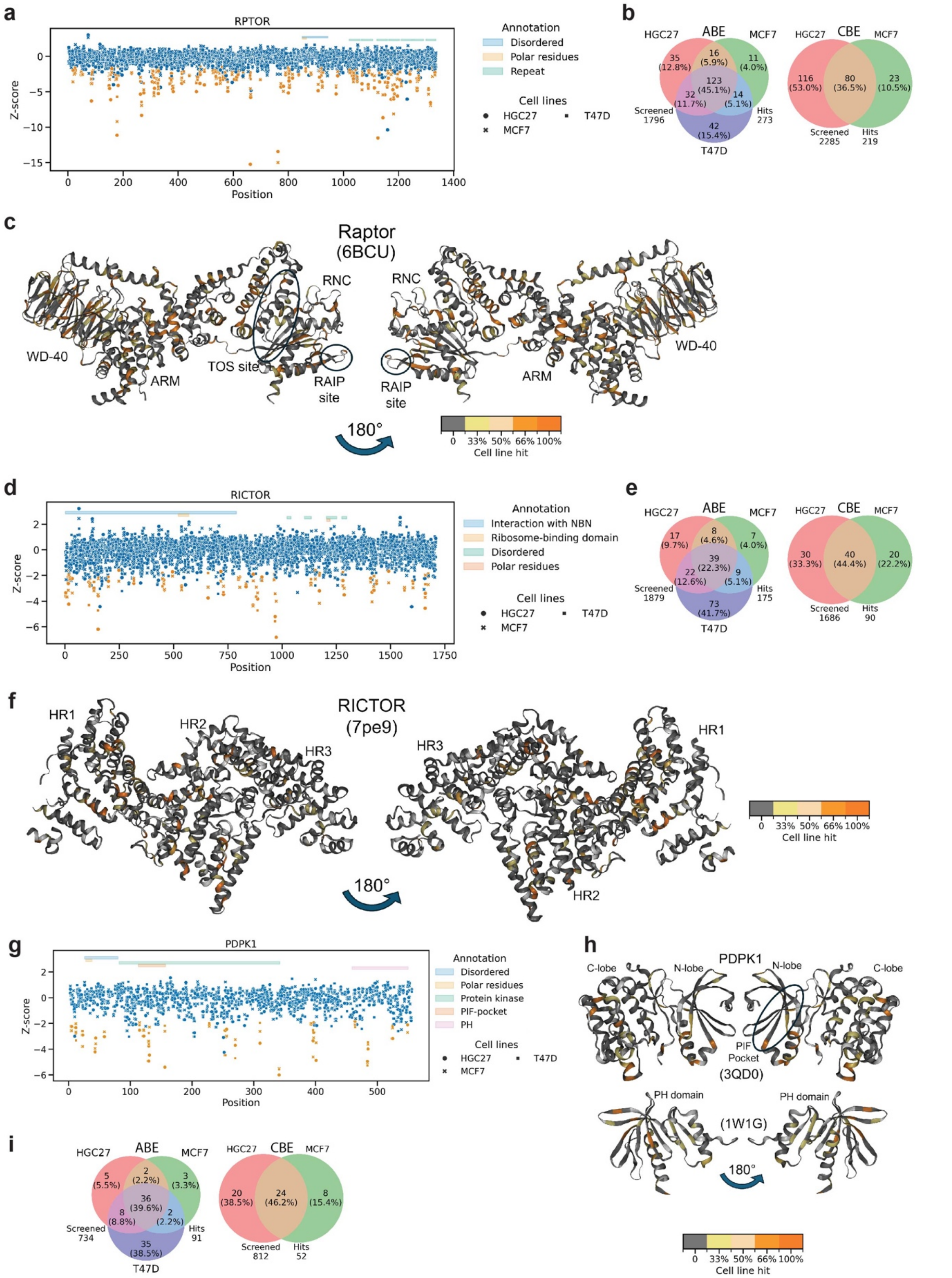


**Supplementary Figure 5: Projection of base editing screen hits onto Raptor, RICTOR and PDPK1 protein 3D structures.** (**a**, **d**, **g**) Z-score for each missense-inducing guide reported on the amino acid residue position for essential genes (a) Raptor, (d) RICTOR and (g) PDPK1. Points coloured in orange highlight guides that are a hit in all three tested cell lines in ABE, or a hit in both HGC27 and MCF7 in CBE. (**b**, **e**, **i**) Venn diagram of the percentage of hits shared between HGC27, MCF7 and T47D cell lines in ABE screens (top) and between HGC27 and MCF7 cell lines in CBE screens (bottom) for (b) *RPTOR,* (e) *RICTOR and* (i) *PDPK1.* (c, f, h) Percentage of cell line hits for each residue shown in the protein structure for (c) Raptor, (f) RICTOR and (h) PDPK1. Raptor structure from PDB code 6BCU, mTOR structure from PDB code 7PE9, and PDPK1 from PDB codes 3QD0 and 1W1G.

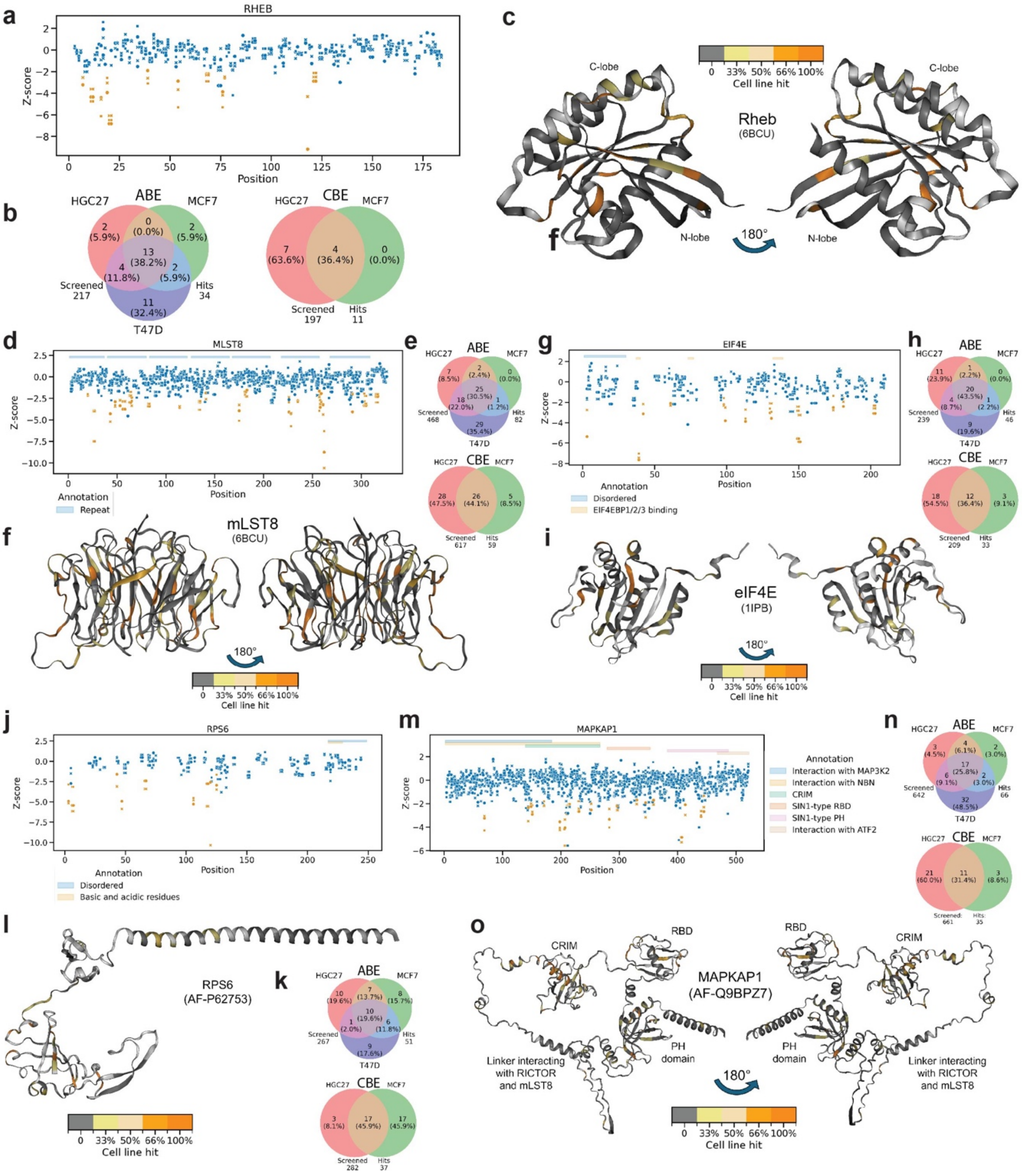


**Supplementary Figure 6: Projection of base editing screens onto RHEB, mLST8, eIF4E, RPS6 and MAPKAP1 protein 3D structures.** (**a, d, g, j, m**) Z-score for each missense-inducing guide reported on the residue position for essential genes (a) RHEB, (d) mLST8, (g) eIF4E, (j) RSP6 and (m) MAPKAP1. Points coloured in orange highlight guides that hit in all three tested cell lines in ABE, or hit in both HGC27 and MCF7 in CBE. HGC27 hits are circles, T47D hits squares and MCF7 hits are depicted as crosses. (**b, e, h, k, n**) Venn diagram of the percentage of hits shared between HGC27, MCF7 and T47D cell lines in ABE screens (top) and between HGC27 and MCF7 cell lines in CBE screens (bottom) for (b) *RHEB*, (e) *MLST8*, (h) *EIF4E*, (k) *RSP6* and (n) *MAPKAP1.* (**c, f, i, l, o**) Percentage of cell line hits for each residue shown in the protein structure for (c) RHEB, (f) mLST8, (i) eIF4E, (l) RPS6 and (o) MAPKAP1. RHEB structure from PDB code 6BCU, mLST8 structure from PDB code 6BCU, eIF4E structure from PDB code 1IPB, RPS6 structure from AlphaFold Database with UniprotID P62753, and MAPKAP1 structure from AlphaFold Database with UniprotID Q9BP27.

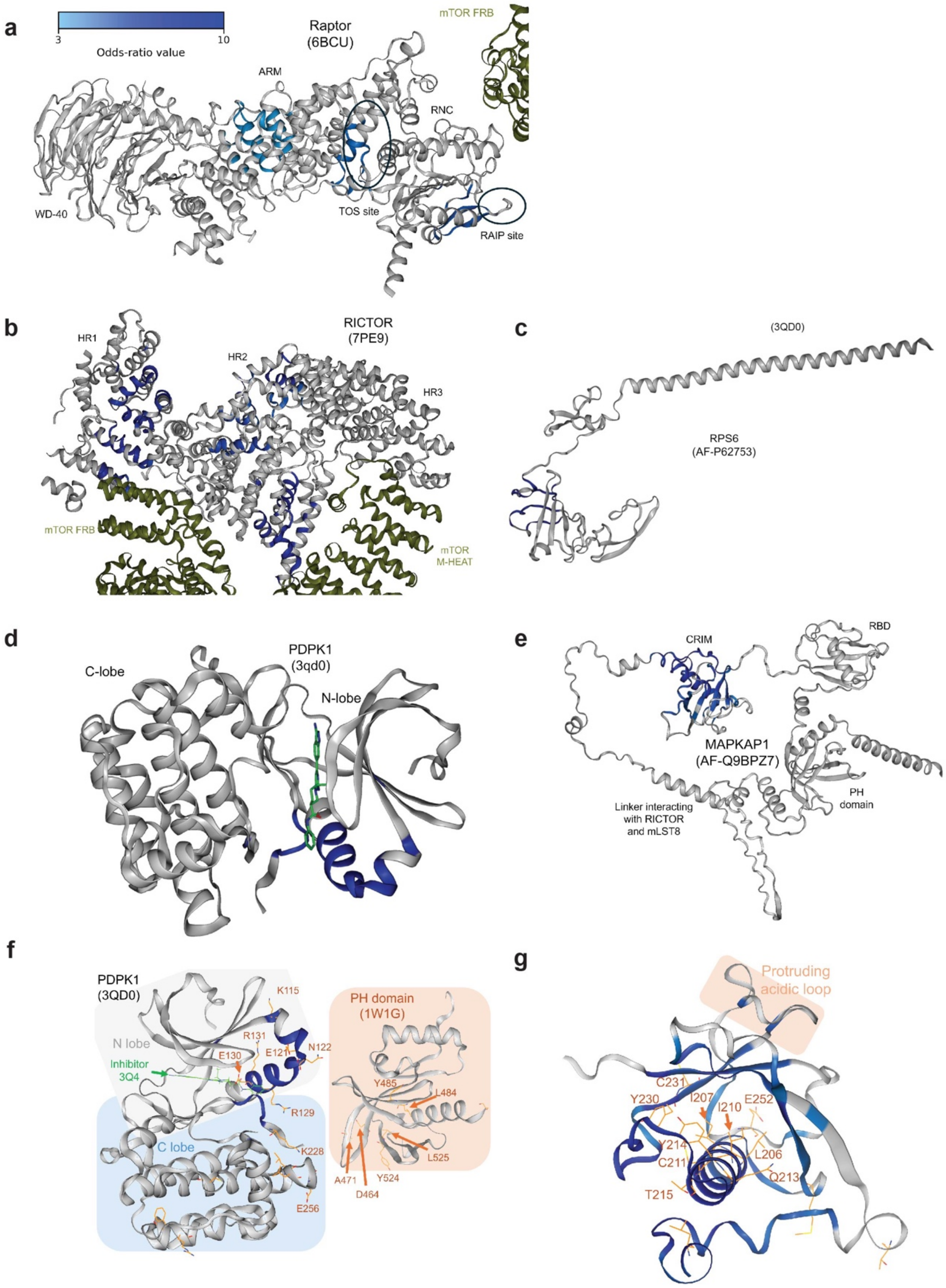


**Supplementary Figure 7: Odds ratio framework for identification of patches within Raptor, RICTOR, PDPK1, RPS6 and MAPKAP1 protein structures.** Significant odds ratio patches on (**a**) Raptor, (**b**) RICTOR, (**c**) RPS6 (**d**) PDPK1 kinase domain and (**e**) MAPKAP1.

The odds ratio patches are coloured with a light blue to dark blue scale for ascending values from 3 to 10. (**f**) Focus on the PDPK1 charged residue hits on the kinase site (PDB code: 3QD0) at the putative interface with the plastron homology (PH) domain (PDB code: 1W1G). (**g**) Cartoon representation of AlphaFold database structure of MAPKAP1 coloured by OR (focused on the CRIM from AlphaFold database structure Q9BPZ7).

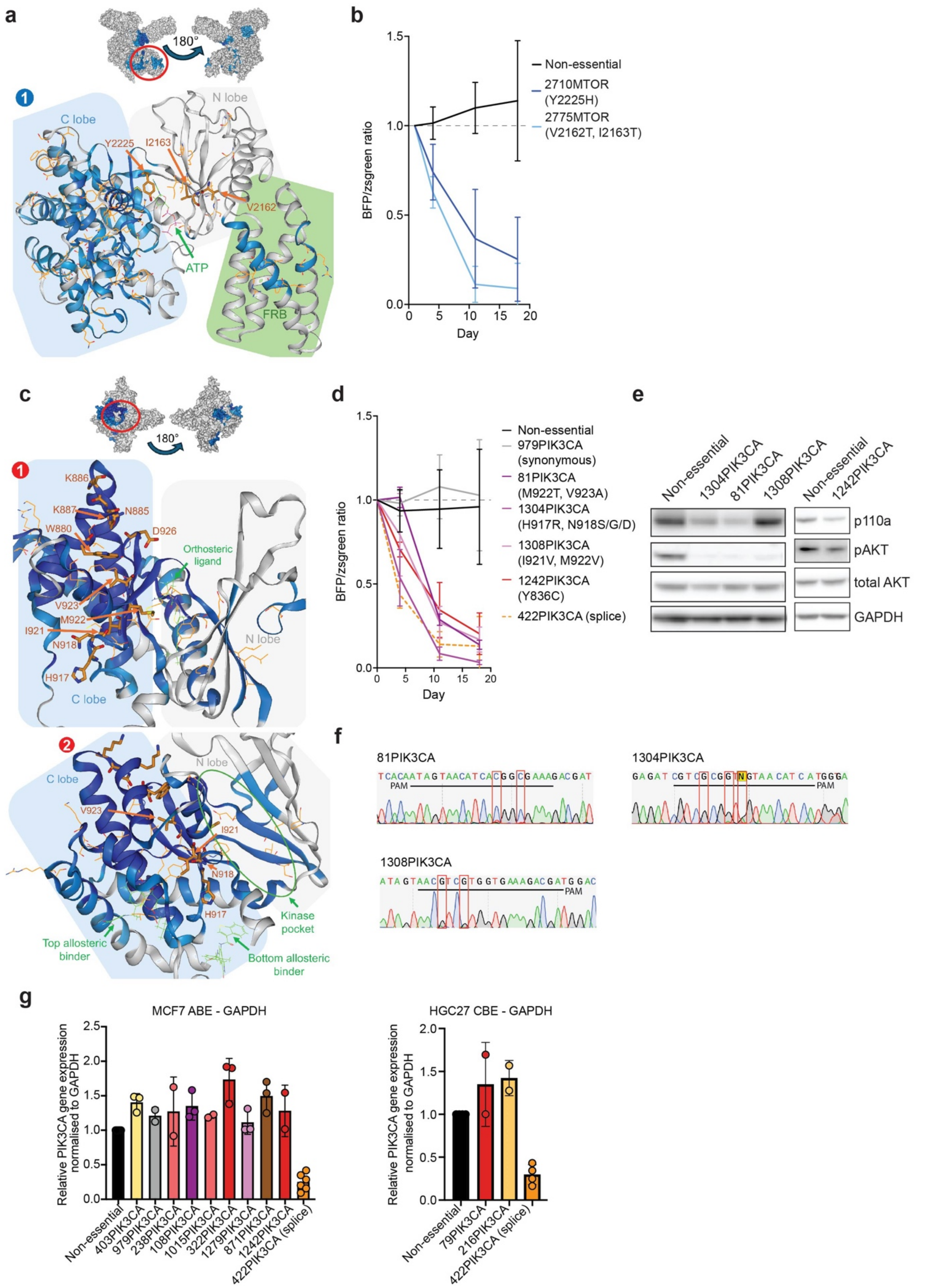


**Supplementary Figure 8: Structural analysis of odds ratio patches on kinase sites on mTOR and p110α. a**) Focus on the kinase orthosteric site of mTOR (PDB code: 6BCU), corresponding to identified mTOR site 1. **b**) Co-competition experiment of non-essential and sgRNAs targeting the mTOR kinase site in MCF7 ABE cells (N=3). **c**) Focus on the kinase orthosteric site (top) and allosteric binding sites (bottom) of p110α close to the C-lobe (PDB code: 8V8J), corresponding to identified p110α sites 1 and 2. **d**) Co-competition experiment of non-essential and sgRNAs targeting the p110α kinase site in MCF7 ABE cells. N=3 for 979PIK3CA and 81PIK3CA, N=5 for 1304PIK3CA and 422PIK3CA, N=4 for 1308PIK3CA and N=2 for 1242PIK3CA. **e**) Western blot of validated sgRNAs compared with non-essential guides in MCF7 ABE cells 7 days after transfection. **f**) Sanger sequencing of sgRNA transfected MCF7 ABE cells showing editing of targeted sites. **g**) qPCR results of edited MCF7 ABE or HGC27 CBE cells 7 days after transfection, *PIK3CA* expression were normalised to *GAPDH* expression in control cells transfected with non-essential sgRNA. N=2–6 biological replicates (dots), bars represent mean ±SD.
